# RbohF-mediated ROS production is required for tapetal PCD and pollen maturation in rice

**DOI:** 10.64898/2026.04.12.718083

**Authors:** Hsiang Chia Lu, Min-Jeng Li, Jo-Chi Hung, Hung-Chien Hsing, Ting-ting Yang, Yi-Ping Ho, Swee-Suak Ko

## Abstract

Reactive oxygen species (ROS) function as critical signaling molecules during plant reproductive development, particularly in regulating tapetal programmed cell death (PCD). However, the molecular mechanisms controlling ROS production in rice anthers remain incompletely understood. In this study, we characterized the role of *RbohF*, a rice respiratory burst oxidase homolog, using CRISPR/Cas9 knockout lines. Loss of *RbohF* resulted in severe defects in anther development, including impaired pollen maturation and reduced fertility. Histological analyses revealed progressive degeneration of microspores and abnormal tapetal persistence in the mutant. ROS staining demonstrated that superoxide and hydrogen peroxide accumulation was significantly reduced and delayed in *rbohf* anthers, particularly at the young microspore stage. Consistently, TUNEL assays showed delayed tapetal PCD in the mutant, indicating that *RbohF*-mediated ROS production is required for the timely initiation of tapetal degeneration. Gene expression analyses further suggested that *RbohF* functions downstream of major transcriptional regulators of pollen development. Notably, the male-sterile phenotype of *rbohf* mutants was exacerbated at cooler temperatures, highlighting a role for RbohF in buffering environmental stress. Together, these findings establish RbohF as a key regulator of ROS homeostasis, integrating developmental and environmental signals to ensure proper pollen development in rice.

## Introduction

Successful pollen development is essential for plant reproduction and crop production. It occurs within the anther and requires coordinated differentiation and a tightly regulated biological process involving complex genetic and physiological networks that govern cell differentiation, metabolic activity, and programmed cell death (PCD). Timely degeneration of the tapetum through PCD is essential for normal pollen maturation and pollen fertility, and disruption of this process frequently results in pollen abortion and male sterility in various plant species, including *Arabidopsis thaliana* and *Oryza sativa* (Bai *et al*., 2019; Feng *et al*., 2012; Ko *et al*., 2014; Li *et al*., 2006; Parish and Li, 2010; Yi *et al*., 2016). Tapetal cells develop alongside microspores and are essential to microsporogenesis. After meiosis, they provide vital proteins, lipids, and structural components—initially through active secretion and eventually via programmed degradation (Parish and Li, 2010; Varnier *et al*., 2005). Tapetal cell degeneration occurs at distinct stages in different plants. For instance, in rice, tapetal degeneration is detected as early as the tetrad stage; whereas in Arabidopsis, visible tapetal degeneration occurs during pollen mitotic division (Uzair *et al*., 2020; Xie *et al*., 2014).

Properly timed tapetal PCD is primarily triggered by peaks in the levels of reactive oxygen species (ROS), such as hydrogen peroxide (H_2_O_2_), the hydroxyl radical (^•^OH), and the superoxide anion radical (O^•^_2_^-^) (Halliwell, 2006). While ROS are potentially hazardous metabolic byproducts capable of inflicting oxidative damage, they simultaneously function as indispensable signaling mediators (Schieber and Chandel, 2014; Sies and Jones, 2020). In rice, several bHLH TFs have been identified as key regulators of tapetal development and degeneration. For example, *Undeveloped Tapetum 1* (*UDT1*/bHLH164) (Jung *et al*., 2005), *Tapetum Degeneration Retardation* (*TDR*/bHLH5) (Li *et al*., 2006), *TDR-Interacting Protein 2* (*TIP2*/bHLH142) (Fu *et al*., 2014; Ko *et al*., 2014), and *Eternal Tapetum 1* (*EAT1*/DTD1/bHLH141) (Ji *et al*., 2013; Niu *et al*., 2013). In addition to bHLH TFs, MYB-type TFs such as *GAMYB* (Aya *et al*., 2009; Liu *et al*., 2010), *MYB80/MYB103* (Lei *et al*., 2022; Phan *et al*., 2011), and *MYB35/TDF1* (Cai *et al*., 2015), also play essential roles in regulating tapetal development and degeneration. These transcription factors form a hierarchical regulatory network that orchestrates gene expression during pollen development, and mutations in these regulators lead to complete male sterility.

In the context of tapetal PCD, NADPH oxidases—also known as respiratory burst oxidase (Rboh) homologs—are major enzymatic sources of ROS production in plant cells (Jimenez-Quesada *et al*., 2016; Xie *et al*., 2014). In Arabidopsis, the NADPH oxidase, *AtRbohE*, has been shown to exhibit a distinct spatiotemporal expression pattern, being primarily localized in the tapetum from stages 6 to 11 of anther development. Consequently, ROS production in the anther is not a continuous process but is governed by a strictly regulated spatiotemporal program (Xie *et al*., 2022; Xie *et al*., 2014). Research on tobacco and tomato further validated the functional conservation of *Rboh* gene across these plant systems (Yu *et al*., 2017). In rice, nine *Rboh* genes have been identified in the genome (Zhu *et al*., 2024). However, the specific *Rboh* genes involved in this process in rice (*Oryza sativa*) have yet to be identified. However, despite the well-established importance of ROS signaling in tapetal PCD, the specific Rboh members responsible for ROS generation during rice anther development remain largely unknown. Identifying these genes is essential for understanding the molecular mechanisms underlying ROS-mediated regulation of pollen development.

In this study, we identified *RbohF* as a gene preferentially expressed in rice anthers. Functional characterization of *RbohF* using CRISPR/Cas9 knockout mutants revealed that disruption of RbohF results in significantly reduced pollen fertility. Through a combination of molecular, histological, and physiological analyses, we demonstrate that RbohF-mediated ROS production is required for proper tapetal PCD and pollen maturation.

## Material and methods

### Generation of CRISPR RbohF knockout mutants

We generated knockout (KO) mutants of the historical Taiwanese Japonica rice variety, TNG67, using the CRISPR/Cas9 system. The CRISPR/Cas9 constructs were designed and prepared in collaboration with the Plant Technology Core Facility at the Agricultural Biotechnology Research Center (ABRC), Academia Sinica. Two specific guide RNA (gRNA) sequences targeting RbohF (Respiratory burst oxidase homolog F) were selected, and the subsequent rice transformation was conducted by the ABRC Plant Genetic Transformation Core Facility. Three independent CRISPR-edited lines (cF#7, cF#8, and cF#15) were cultivated for phenotypic observation during the 2023-II and 2024-I cropping seasons at the GMO greenhouse at the Biotechnology Center in Southern Taiwan (AS-BCST), located in the Tainan Science Park (23°06’14.4"N, 120°17’31.2"E).

### Histological analysis of anthers

To closely examine the anther development of the *rbohf* mutant, transverse sections of both wild-type (WT) and *rbohf* anthers were prepared. These samples, ranging from the meiotic to the vacuolated pollen stages, were processed using the LR White resin-embedding technique for high-resolution microscopy.

### TUNEL assay for tapetal programmed cell death

To evaluate tapetal programmed cell death (PCD), anthers from wild-type (WT) and *rbohf* mutant plants were collected from the meiotic to the vacuolated pollen stages. Tissues were fixed, embedded in paraffin, and transversely sectioned (10 µm thickness). DNA fragmentation was detected using the DeadEnd™ Fluorometric TUNEL System (Promega) following a previously described protocol (Ko *et al*., 2014).

### Detection of ROS production in anthers

Anthers from wild-type (WT) and *rbohf* mutant plants were collected at various developmental stages, embedded in paraffin, and sectioned. To detect reactive oxygen species (ROS), sections were stained according to the protocol described by Ko *et al*. (2017). Specifically, superoxide anions were visualized using 0.5 mM nitroblue tetrazolium (NBT), and hydrogen peroxide (H_2_O_2_) was detected with 3,3′-diaminobenzidine (DAB).

### In situ hybridization analysis

Anthers from wild-type (WT) and *rbohf* mutant plants were collected at multiple developmental stages. Tissues were fixed, embedded in paraffin, and transversely sectioned at a thickness of 10 µm. *In Situ* Hybridization (ISH) analysis was subsequently performed using DIG-labeled RNA probes to visualize the localization of *RbohF* transcripts.

### mRNA library construction and sequencing

One mircogram of total RNA was used for library preparation. For mRNA enrichment, poly(A) mRNA was isolated using Oligo(dT) beads; alternatively, ribosomal RNA (rRNA) was depleted to enrich for both coding and non-polyadenylated transcripts, depending on sample quality. The mRNA was then fragmented using divalent cations at high temperatures. First-strand cDNA was synthesized using reverse transcriptase and random primers, followed by second-strand synthesis. To construct strand-specific libraries, dTTP was replaced with dUTP during second-strand synthesis, and the resulting dUTP-labeled strands were selectively degraded using the USER enzyme prior to PCR amplification. The purified double-stranded cDNA underwent end-repair and dA-tailing in a single reaction, followed by T-A ligation of adaptors. Size selection was performed using DNA Clean Beads. Finally, the adaptor-ligated products were amplified using P5 and P7 primers. The validated libraries were multiplexed and sequenced on an Illumina NovaSeq platform in a 2 × 150 bp paired-end (PE) configuration.

### RNA-seq data analysis

Raw sequencing reads in FASTQ format were processed using Cutadapt (v.1.9.1) to remove technical sequences and low-quality bases. Clean reads were aligned to the reference genome (*Oryza sativa*, assembly version) using HISAT2 (v.2.2.1), a splice-aware aligner. The reference genome and annotation files were obtained from ENSEMBL, NCBI, or other databases. Gene-level counts were quantified using HTSeq (v.0.6.1). Reads overlapping exons were counted, and multi-mapping reads or reads overlapping multiple genes were excluded. Differentially expressed genes (DEGs) were identified using DESeq2 (v.1.34.0), which employs a negative binomial distribution model.

### Statistical analysis

Statistical analyses were performed to evaluate differences between the wild-type and CRISPR mutant groups. Comparisons between the two groups were conducted using Student’s t-test (two-tailed). A P value of less than 0.05 was considered statistically significant.

## Results

### RbohF as a putative executor of ROS-dependent programmed cell death during rice anther development

Previously, we found that the *bHLH142* mutant exhibited defective tapetal programmed cell death (PCD) and failed to produce viable pollen (Ko *et al*., 2014). Transcriptome analysis of the *bHLH142* mutant revealed that *RbohF* (Os08g0453700) was significantly downregulated, suggesting that it may function as a downstream component of the bHLH142-mediated regulatory network. Given that bHLH142 plays a key role in regulating tapetal PCD and maintaining reactive oxygen species (ROS) homeostasis during pollen development, we hypothesized that RbohF may function as a ROS-generating enzyme within this pathway. To investigate the potential role of RbohF in rice reproductive development, we first examined its expression profiles using publicly available transcriptome datasets. Data obtained from the RiceXPro database and the Rice eFP Browser indicated that *RbohF* is predominantly expressed in anthers (∼0.7 mm in length) and in panicles during developmental stages P3 to P5 (**Supplementary Fig. S1**). We further validated the expression pattern of *RbohF* in the TNG67 cultivar using reverse transcription quantitative PCR (RT-qPCR). Our results demonstrated that *RbohF* is highly expressed in 9-cm panicles and during the meiotic stage of pollen development (**Fig. 1A,B**).

**Fig. 1.**
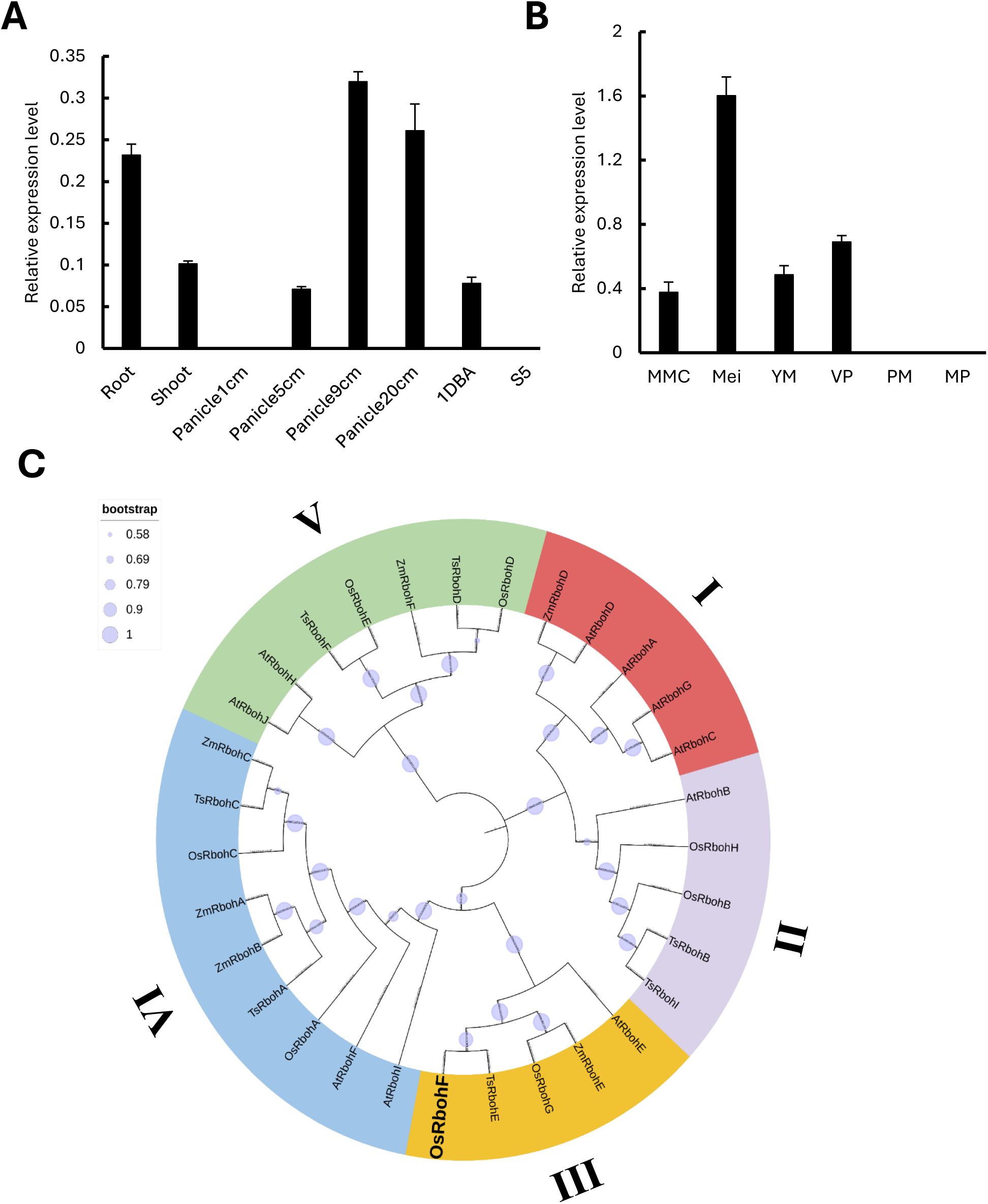
Gene expression patterns and phylogenetic relationships of RbohF. (A) Expression pattern of *RbohF* in various tissues of the TNG67. 1 DBA, one day before anthesis. S5, mature seed. (B) Expression pattern of *RbohF* at various anther developmental stages. MMC, microspore mother cell; Mei, meiosis; YM, young microspore; VP, vacuolated pollen; PM, pollen mitosis; MP, mature pollen. (C) Phylogenetic relationships among plant RBOH family members. Amino acid sequences Rboh from *Arabidopsis*, *Oryza sativa*, *Triticum aestivum*, and *Zea mays* were aligned by ClustalW under default settings. A phylogenetic tree was constructed using neighbor-joining (NJ) with the JTT model method in MEGA 12. Bootstrap support values from 1000 replicates are indicated by blue dots of varying widths, with the width corresponding to the value. Five subgroups are indicated with I–V, respectively. The OsRbohF gene characterized in this study is highlighted in bold.

To explore the evolutionary relationship between RbohF and other plant NADPH oxidases, we conducted a phylogenetic analysis using Rboh protein sequences from rice and other plant species. The resulting phylogenetic tree showed that *RbohF* clusters within subgroup III, together with *OsRbohG*, *TsRbohE*, *AtRbohE*, *ZmRbohE*, and *ZmRbohF* (**Fig. 1C**). Notably, *AtRbohE* has been reported to be specifically expressed in the tapetum and is required for ROS production and proper pollen development in Arabidopsis (Xie *et al*., 2014). The close phylogenetic relationship between *OsRbohF* and *AtRbohE* therefore suggests that these proteins may share conserved biological functions.

### Targeted mutagenesis of RbohF using CRISPR/Cas9 in TNG67 rice

To further investigate the functional role of *RbohF* in rice pollen development, we generated knockout mutants in the TNG67 background using CRISPR/Cas9 technology. Two single-guide RNA (sgRNA) target sites were strategically designed within exons 6 and 8 of the *RbohF* gene (**Fig. 2A**). Among 22 independent T_1_ transgenic lines, three representative mutant alleles—*rbohf*-7-2, *rbohf*-8-1, and *rbohf*-15-7— were identified, each exhibiting distinct mutation patterns (**Fig. 2A**). All three lines contained mutations at both target sites, resulting in premature termination codons that are predicted to produce truncated, non-functional RbohF protein (**Fig. 2B, C**). Genotype verification via genomic PCR confirmed that the T_2_ progeny carried the expected mutations. Specifically, lines *rbohf*-7-2-1, *rbohf*-7-2-2, *rbohf*-8-1-25, and several T_2_ individuals from *rbohf*-15-7 were identified as homozygous mutants that had segregated away the hygromycin resistance marker (**Supplementary Fig. S2**).

**Fig. 2.**
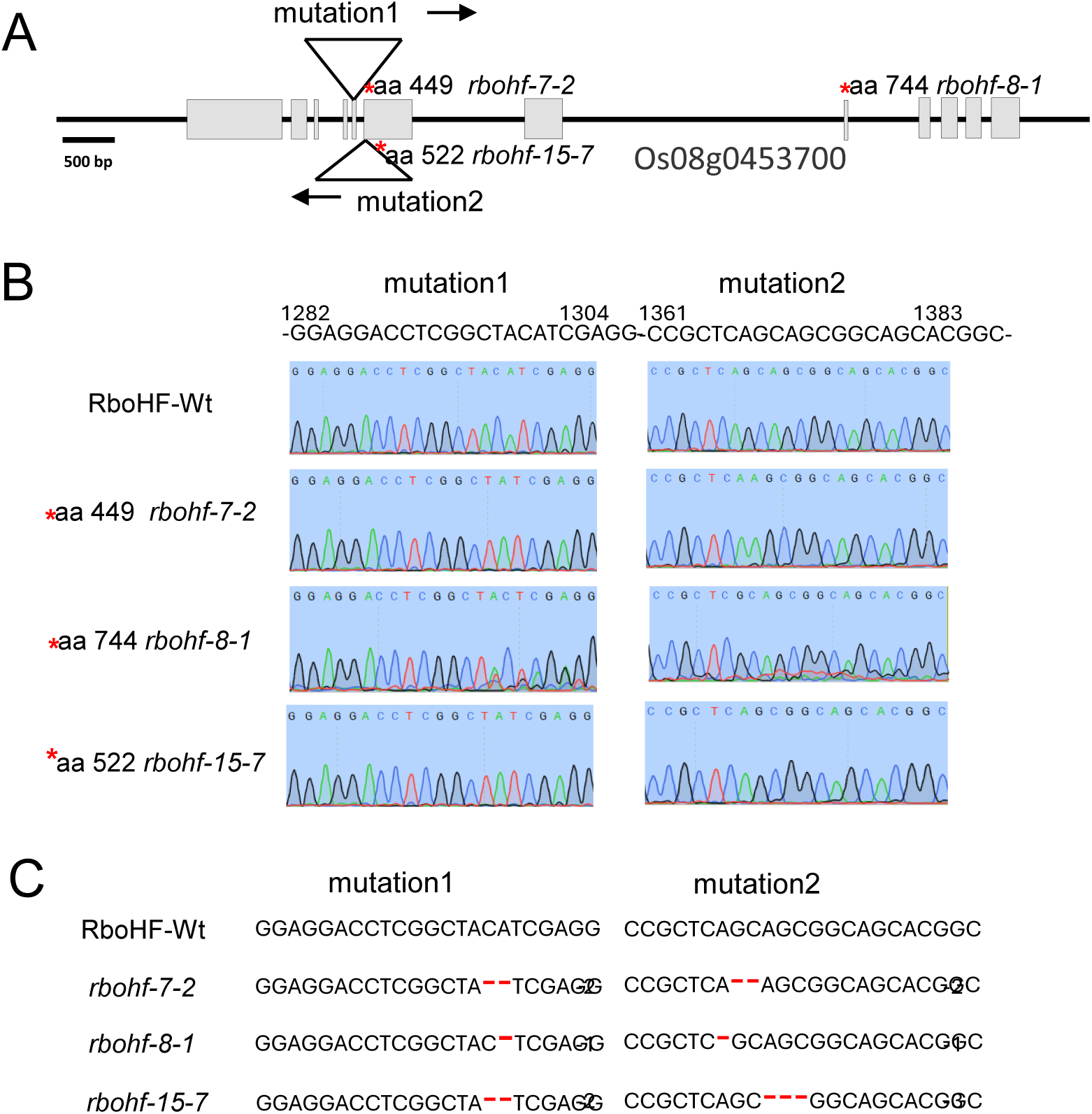
Gene editing *RbohF* using the CRISPR Cas9 system. (A) A schematic diagram of the *RbohF* gene structure and sgRNA target sites. The gray boxes and lines represent exons and introns, respectively. Scale bar: 500 bp. Two sgRNAs, designated as mutation 1 and mutation 2, were designed to target the third and fourth exons. (B) Gene sequencing results of CRISPR *rbohf* T_2_ transgenic lines. *, site of stop codon in the *rbohf* mutants. (C) Identified mutant genotypes in the three CRISPR *rbohf* transgenic lines. The sequences highlight the specific deletions (indicated by red dashes) at both target sites.

### Disruption of RbohF reduced pollen fertility

The *rbohf* CRISPR-knockout lines were characterized over two consecutive cropping seasons (2023-II and 2024-I) in the GMO net house at the Biotechnology Center in Southern Taiwan (AS-BCST). All mutant lines exhibited normal vegetative development consistent with the TNG67 wild-type (WT) under both seasonal conditions (**Fig. 3A, 4A**). However, striking differences were observed in reproductive organs. During the 2024-I (Spring) season, mutant lines displayed distinct anther phenotypes, characterized by smaller dimensions and a markedly paler yellow coloration relative to the WT (**Figs. 3B and 4B**). Iodine-potassium iodide (I_2_-KI) staining confirmed significant reductions in pollen viability in all CRISPR lines across both seasons (**Fig. 3C, 4C**). In the 2023-II season, the percentage of fully stained (viable) pollen in lines *rbohf*-7, *rbohf*-8, and *rbohf*-15 was approximately 30%, 42%, and 65%, respectively, significantly lower than the WT (**Fig. 3C**). A similar trend was recorded during the 2024-I season, with viability values further declining to approximately 20%, 40%, and 60%, respectively (**Fig. 4C**). Consequently, the grain fertility of all three *rbohf* knockout lines remained significantly lower than that of the WT in both cropping seasons (**Fig. 3D, E** and **4D, E**). Temperature records (max./mean/min.) during rice pollen developmental stages were 30.9/24.7/20.6°C in the second cropping season of 2023, whereas those in the first cropping season of 2024 were lower at 26.6/20.1/15.7°C (**Supplementary Fig. S3**).

**Fig. 3.**
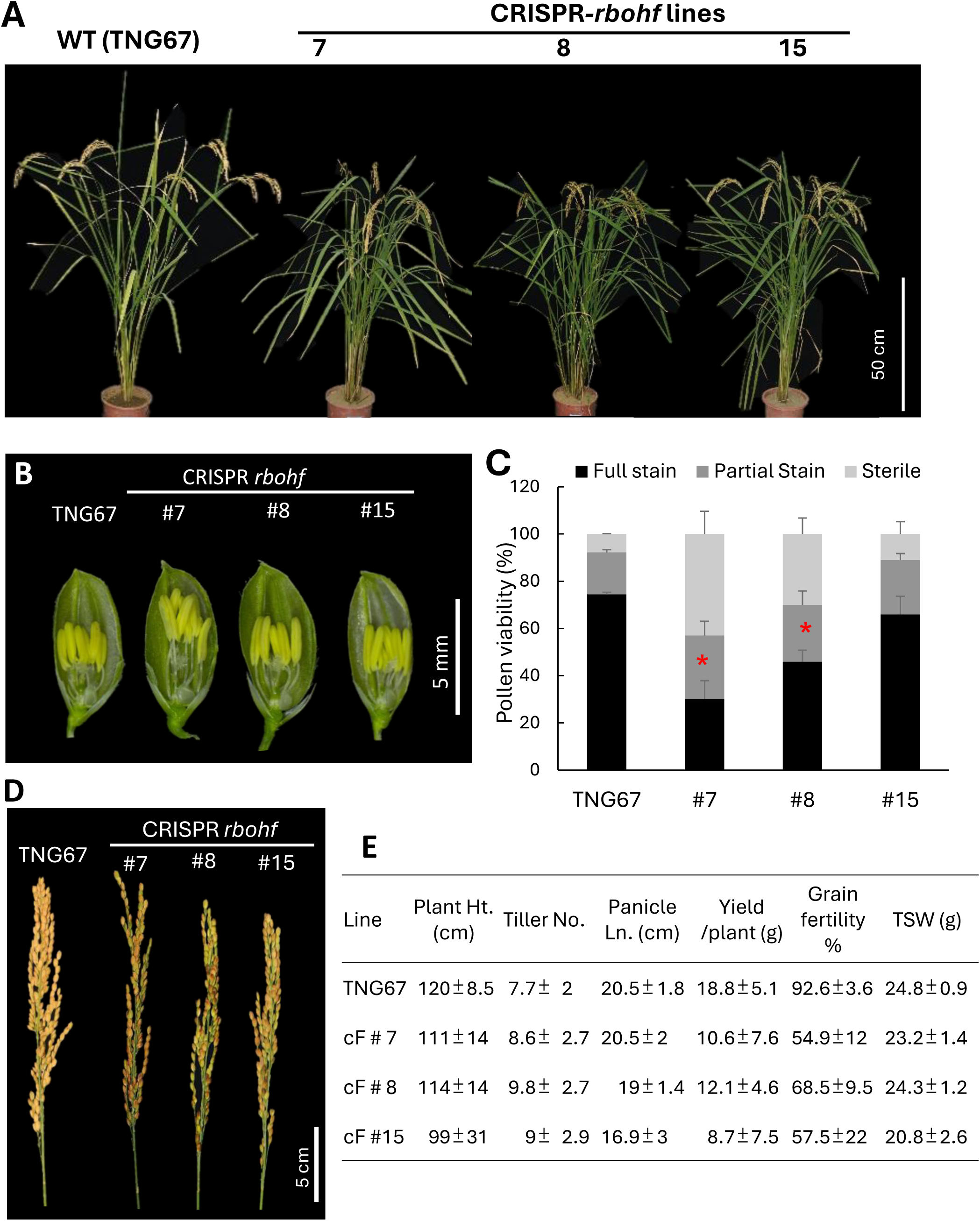
Agronomic traits and pollen fertility of wild-type TNG67 and CRISPR *rbohf* T_3_ lines during the 2023-II cropping season. (A) Plant morphology at maturity. (B) Spikelets at one day before anthesis. Scale bar: 5 mm. (C) Pollen viability of WT and three CRISPR lines. *, significant differences between WT and CRISPR lines of the full stained pollen were determined using Student’s t-test at *p* < 0.05. (D) Panicles at the mature stage. Scale bar: 5 cm. (E) Yield and agronomic traits (n= 12 plants).

**Fig. 4.**
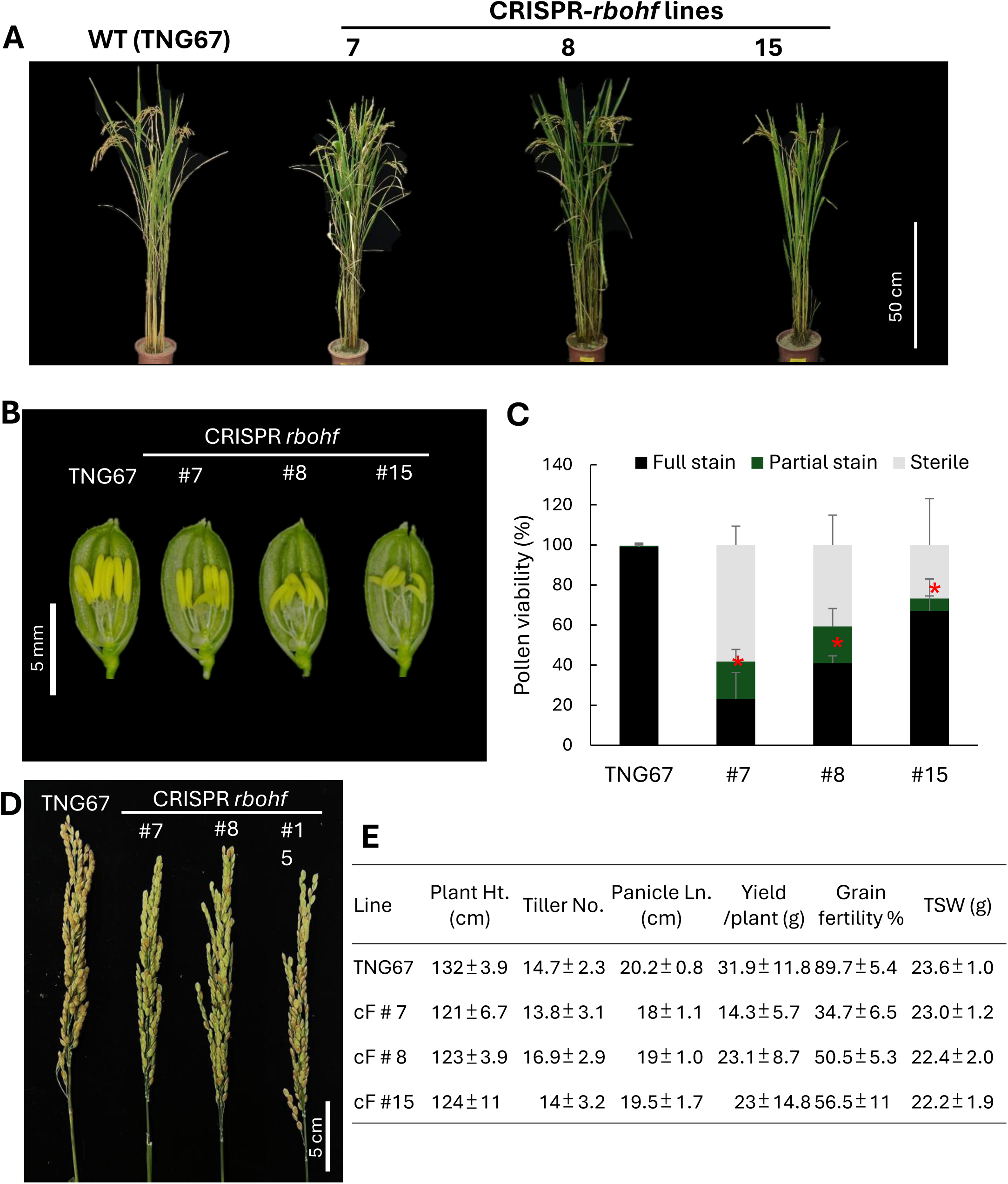
Agronomic traits and pollen fertility of wild-type TNG67 and CRISPR *rbohf* T_4_ lines during the 2024-I cropping season. (A) Plant morphology at maturity. (B) Spikelets at one day before anthesis. Scale bar: 5 mm. (C) Pollen viability of WT and three CRISPR lines. *, significant differences between WT and CRISPR lines of the full stained pollen were determined using Student’s t-test at *p* < 0.05. (D) Panicles at the mature stage. Scale bar: 5 cm. (E) Yield and agronomic traits (n = 12 plants).

### Disruption of RbohF impaired anther development

To investigate the cellular basis of pollen abortion in *rbohf* mutants, we conducted a comparative histological analysis of anther sections from the wild-type (TNG67) and mutants across three key developmental stages: meiosis, the microspore stage, and the vacuolated pollen stage (**Fig. 5**). In TNG67, microspore mother cells underwent normal meiotic progression, developing into distinct, well-organized microspores, and subsequently developed into vacuolated pollen. In contrast, *rbohf* mutants exhibited progressive cellular abnormalities throughout anther development. The mutants failed to enter meiosis properly, and early signs of meiocyte degeneration were evident. As development progressed to the microspore stage, degeneration became more pronounced, with numerous microspores appearing collapsed (**Fig. 5**). At the vacuolated pollen stage, mutant anthers contained severely shrunken and malformed pollen grains that lacked organized cytoplasmic contents compared to the wild-type (**Fig. 5**).

**Fig. 5.**
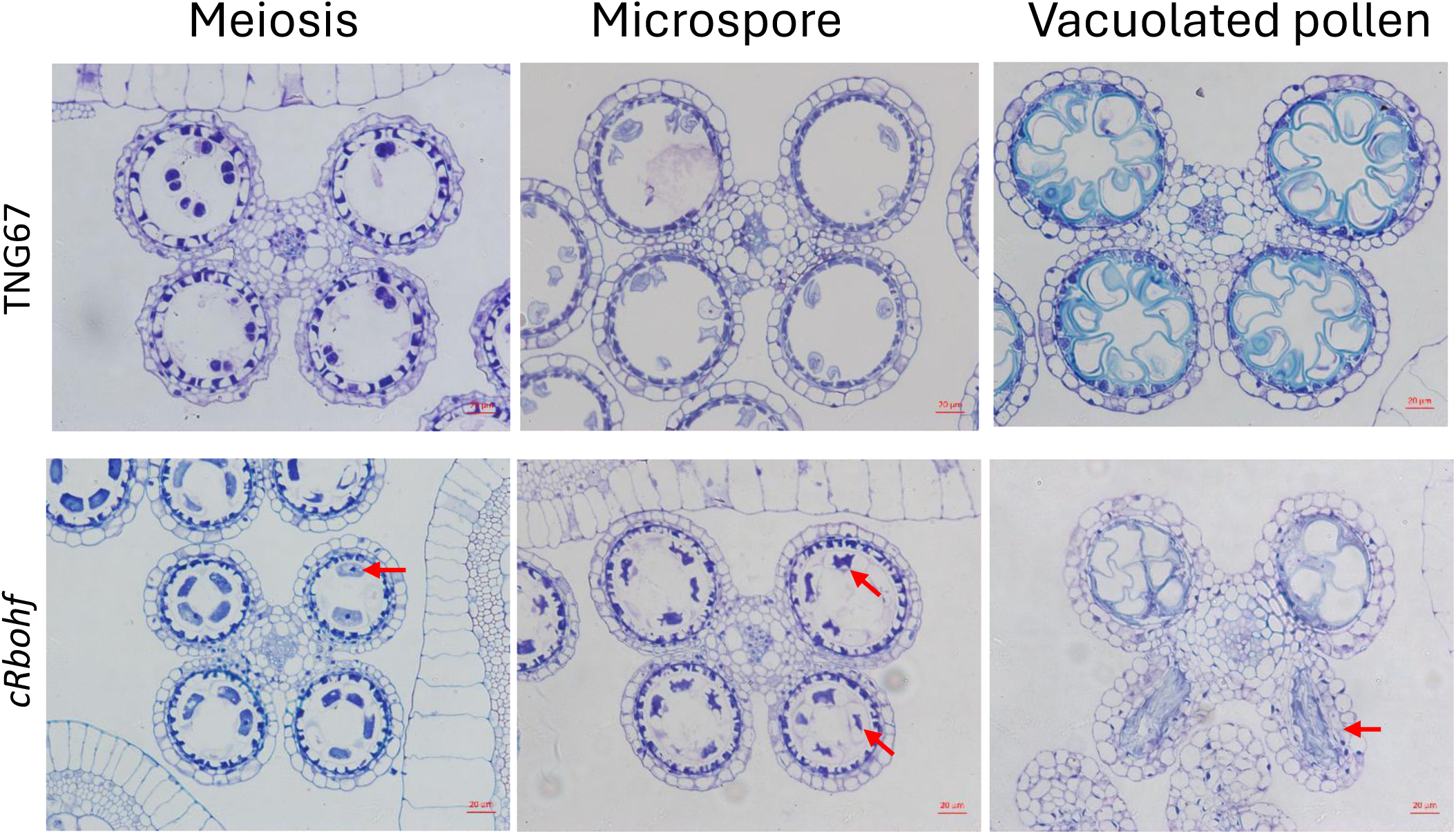
Anther development anatomy in wild-type (TNG67) and CRISPR *rbohf* mutant plants. Representative images showing the developmental progression in wild-type (TNG67) and *rbohf* mutant plants across three stages: meiosis, microspore, and vacuolated pollen. Arrows indicate degenerated microspores and vacuolated pollen observed in the *rbohf* mutant.

## Spatiotemporal expression pattern of RbohF in developing anthers

To further elucidate the spatial and temporal expression profiles of *RbohF* during pollen development, we performed RNA fluorescence *in situ* hybridization (FISH) on transverse anther sections from wild-type (TNG67) and *rbohf* mutant plants. In the WT, *RbohF* transcripts were detected from the meiosis stage onward, with the hybridization signal primarily localized within the tapetal layer and developing microspore mother cells. The gene expression intensity peaked at the microspore stage, with strong, distinct signals detected in both the tapetum and newly released microspores. As development progressed to the vacuolated pollen stage, the *RbohF* signal was no longer detectable (**Fig. 6**). In contrast, little to no hybridization signal was observed in the tapetum of the *rbohf* mutant anthers across all examined developmental stages (**Fig. 6**).

**Fig. 6.**
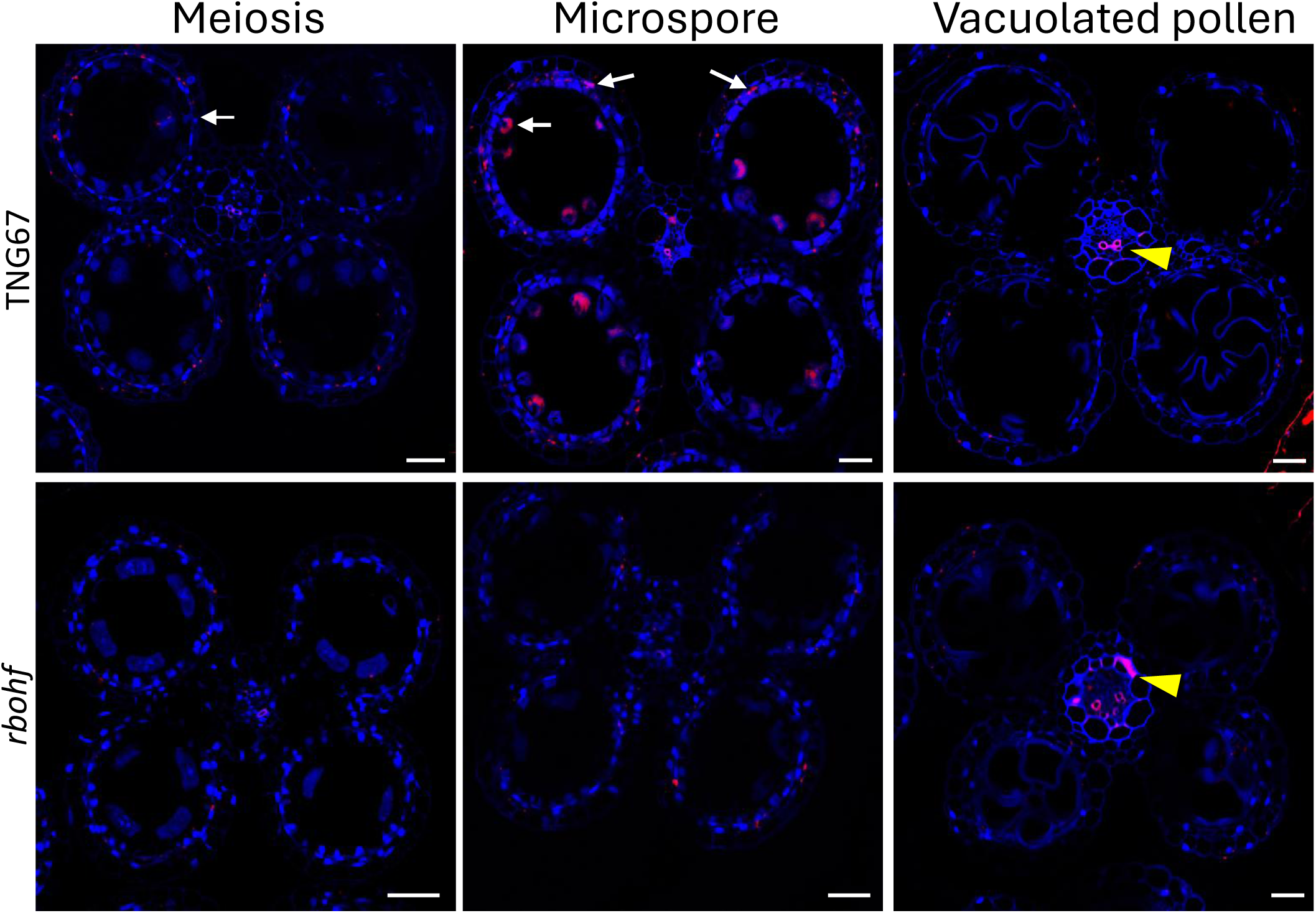
RNA fluorescence *in situ* hybridization (FISH) study of the expression patterns of *RbohF* in the wild-type TNG67 and *rbohf* mutant. Anther tissue embedded in paraffin was transversely sectioned to 10 µm. Another tissue was stained with TSA working solution containing Cyanine 5 Plus Amplification Reagent (300× dilution). Ex/Em: 639/740 nm. DAPI signal was excited by a UV filter at a wavelength of 405 nm. The arrows indicate the gene expression patterns of *RbohF* (red color). Arrowheads show the autofluorescence signal. Tissues were counterstained with DAPI (blue color). Scale bars: 20 µm.

### Loss of RbohF disrupted ROS homeostasis in developing anthers

Given that Rboh proteins are major producers of reactive oxygen species (ROS), we examined the spatial and temporal distribution of superoxide (O^•^_2_^-^) and hydrogen peroxide (H_2_O_2_) in developing anthers using NBT and DAB whole-mount staining of spikelets. In WT anthers, ROS accumulation followed a stage-specific pattern, with superoxide and H_2_O_2_ becoming detectable at the meiosis stage and peaking during the young microspore (YM) stage (**Supplementary Fig. S4 A,C, upper panels**). In contrast, in the anthers of the *rbohf*-7-2 line, the intensity of both O^•^_2_^-^ and H_2_O_2_ signals was significantly reduced at the YM stage compared with WT (**Supplementary Fig. S4 A,C, lower panels**). In addition, a clear delay in the onset of ROS production was observed in the mutant anthers relative to the WT (**Supplementary Fig. S4 B,D, lower panels**).

To examine ROS accumulation at the cellular level in anther tissues, samples were embedded in paraffin, transversely sectioned to 10 µm, dewaxed, and subsequently stained with NBT and DAB to detect superoxide and hydrogen peroxide, respectively. In WT anthers, superoxide accumulated predominantly in the tapetal layer and vascular bundle at the meiosis-II stage and gradually increased at the young microspore (YM) stage (**Fig. 7A**, upper panel). In contrast, the mutant exhibited delayed and weaker superoxide signals at the YM stage (**Fig. 7A**, lower panel). Consistent with the superoxide pattern, H_2_O_2_ accumulation in mutant anthers was also delayed and reduced compared with WT (**Fig. 7B**).

**Fig. 7.**
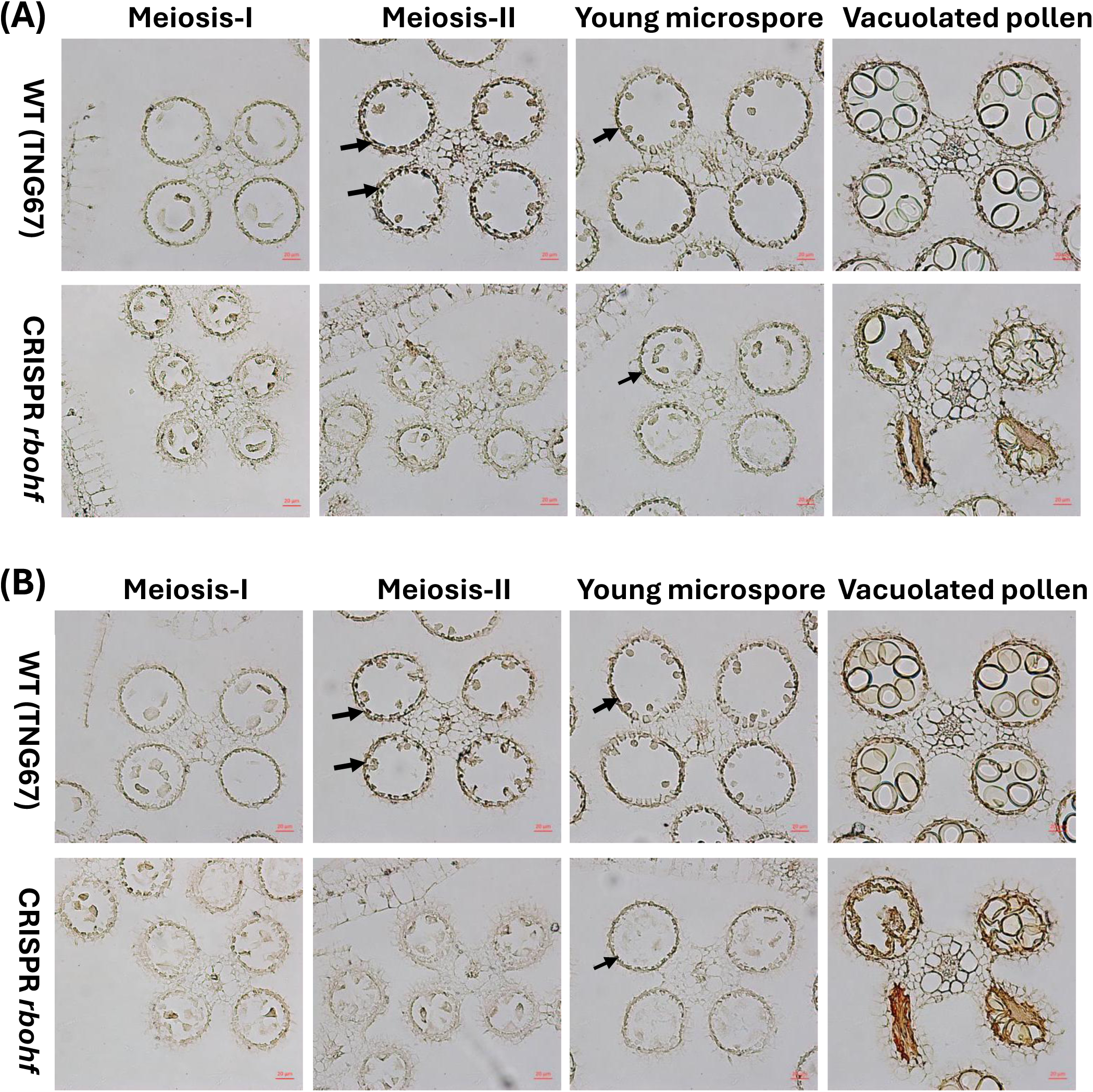
Reactive oxygen species (ROS) staining of anther cross-sections from TNG67 (wild type, WT) and CRISPR/Cas9-mediated *RbohF* knockout mutant during anther development. (A) Detection of superoxide accumulation using nitroblue tetrazolium (NBT) staining. (B) Visualization of H_2_O_2_ using 3,3′-diaminobenzidine (DAB) staining. Arrows indicate ROS signals. Scale bars: 200 µm.

### Delayed tapetal cell PCD in rbohf CRISPR knockout mutant

To assess the progression of tapetal programmed cell death (PCD), a TUNEL assay was performed on anthers from WT and *rbohf* mutants at key developmental stages, from early meiosis to vacuolated pollen. In WT anthers, strong TUNEL signals were detected in tapetal cells at the meiosis stage, indicating timely initiation of PCD (**Fig. 8**, upper panel). The signal intensity subsequently declined as development progressed, consistent with the completion of tapetal degeneration. In contrast, *rbohf* mutant anthers exhibited a clear delay in tapetal PCD. TUNEL signals were weak or barely detectable at the meiosis stage; they only became evident at the YM stage, where the signal remained comparatively weaker (**Fig. 8**, lower panel) than in the WT meiosis stage.

**Fig. 8.**
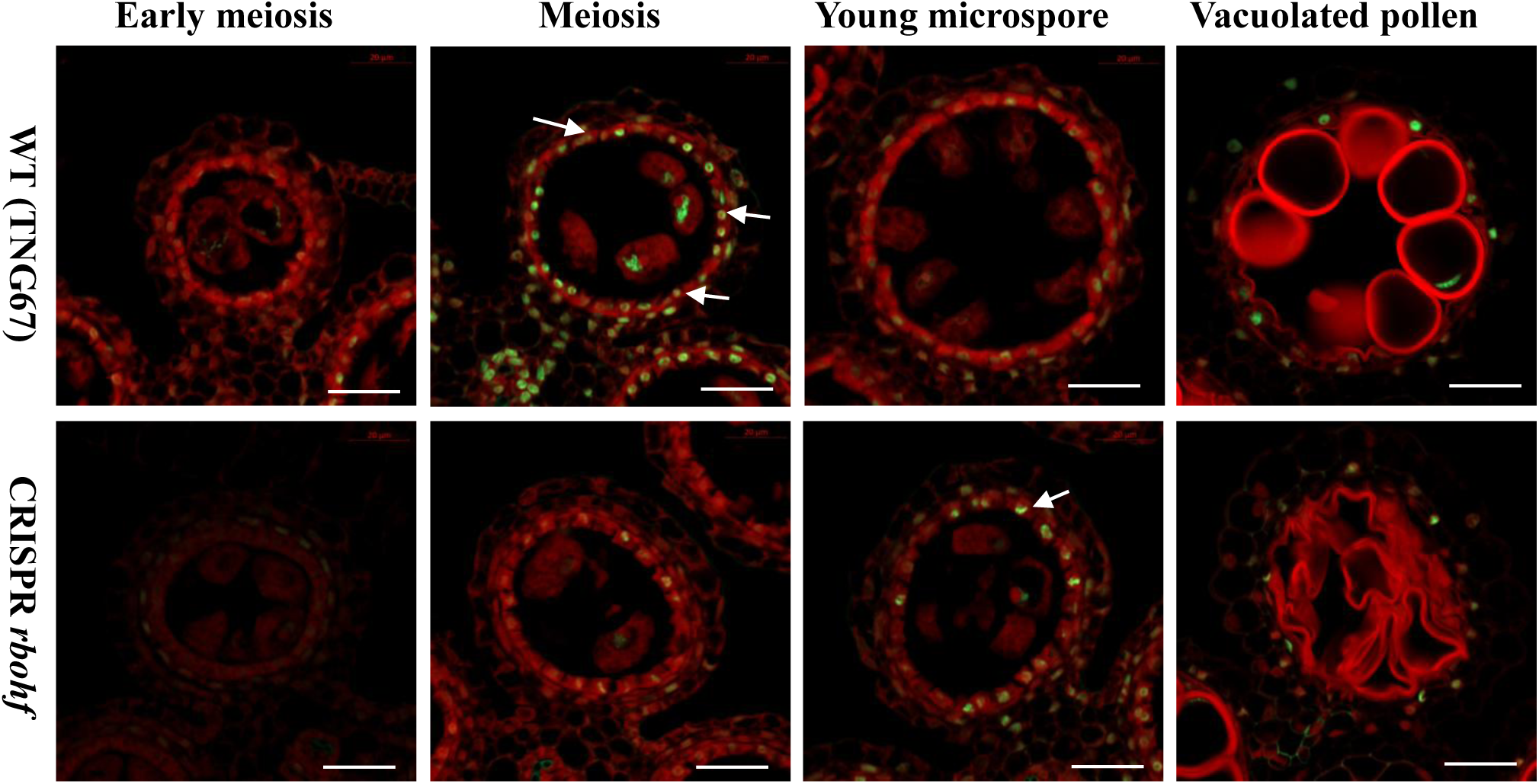
TUNEL assay revealing programmed cell death (PCD) dynamics during anther development in TNG67 (wild type, WT) and CRISPR/Cas9-mediated *RbohF* knockout lines. DNA fragmentation signals (yellow fluorescence). The red signal is propidium iodide staining, and the yellow fluorescence is the merged signal from TUNEL (green) and propidium iodide staining (red). Arrows indicate the tapetal PCD signal. Scale bars: 20 µm.

### RbohF acts downstream of key tapetal transcription factors

To determine the genetic position of RbohF within the regulatory network of rice pollen development, its expression was examined in several well-characterized tapetal transcription factor mutants, including *udt1*, *gamyb-2*, *ms142*, *tdr*, and *eat1*. Gene expression analysis using quantitative RT-PCR performed on anthers at the meiosis stage revealed that the transcript levels of *RbohF* were significantly downregulated in all five homozygous mutant lines compared to the wild-type (**Fig. 9**). Notably, the expression of *RbohF* was nearly abolished in the *gamyb-2*, *ms142*, and *tdr1* mutants, while it was substantially reduced in the *udt1* and *eat1* mutants (**Fig. 9**). These findings demonstrate that gene hierarchy of *RbohF* acts downstream of these key transcription factors.

**Fig. 9.**
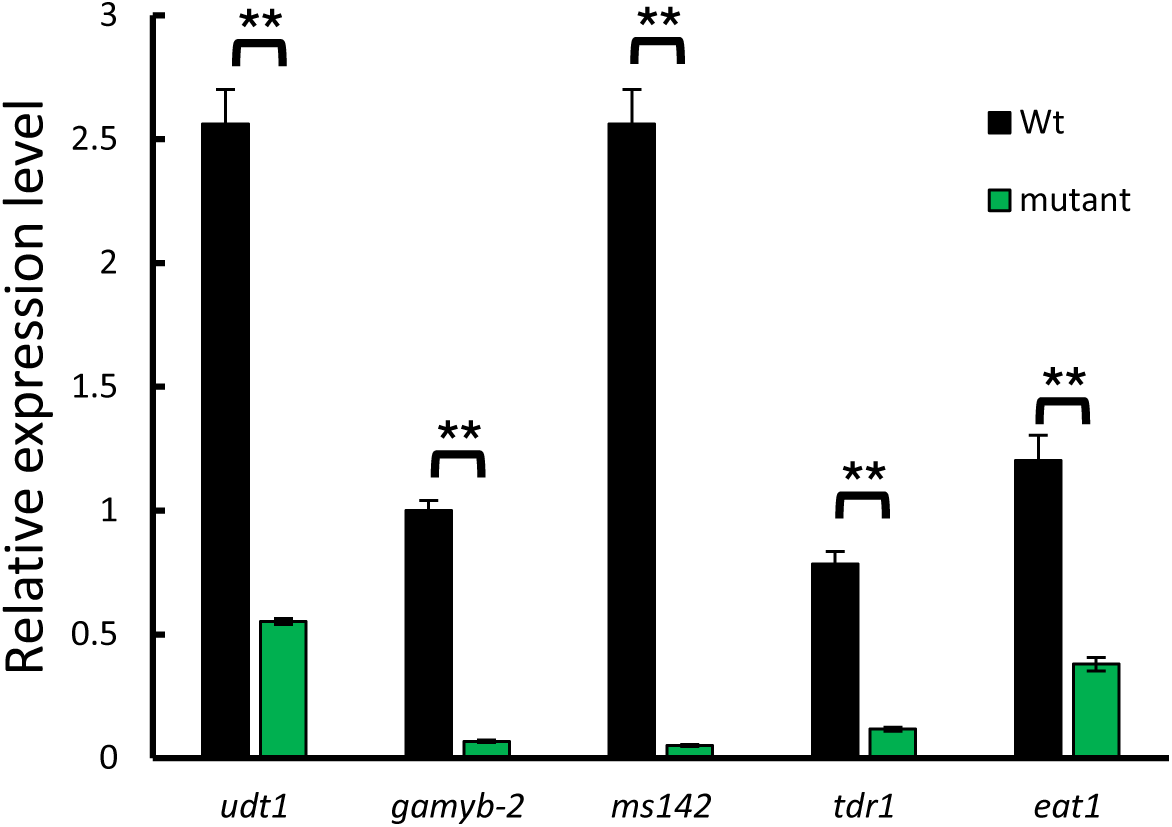
*RbohF* acts downstream of several pollen transcription factors (TFs). Known pollen TF mutants defective in tapetal PCD, including *udt1*, *gamyb-2*, *ms142*, *tdr1*, and *eat1* homozygotes, were used to examine the expression level of *RbohF*. Anthers at the meiosis stage were collected for RNA isolation and RT-qPCR analysis. Statistical significance was determined using Student’s t-test (*p* < 0.01, **).

### Mutant altered gene expression of known pollen markers

To investigate whether the anther developmental defects in the *rbohf* mutant were associated with transcriptional changes in key regulators of pollen development, we performed RT-qPCR analysis of several transcription factors. The expression levels of *UDT1*, *GAMYB*, *TDR*, and *EAT1* did not differ significantly between the WT and *rbohf* knockout lines, suggesting that the disruption of *RbohF* does not affect the upstream transcriptional network controlling these regulators. Notably, *bHLH142* showed a moderate but consistent upregulation in the *rbohf* mutant (**Supplementary Fig. S5**). In addition, several known pollen marker genes associated with PCD revealed that *CP1*, *AP25, AP37, and NADPH* were moderately upregulated in the knockout line at the young microspore stage (YM). In contrast, *C6*, a gene involved in orbicules *(i.e.*, Ubisch bodies) and pollen exine development, was significantly downregulated in the knockout line (**Supplementary Fig. S6**).

### Transcriptome profiling of rbohf mutants during anther development

To elucidate the molecular mechanisms underlying the defective pollen development in *rbohf* mutants, we performed a comparative transcriptomic analysis (RNA-seq) on anthers from the CRISPR-generated *rbohf* lines and the wild-type (WT) at two critical developmental stages: the meiosis (Mei) stage and the young microspore (YM) stage. Differentially expressed genes (DEGs) were identified using a stringency threshold of Fold Change > 2 and an adjusted P value of *p* < 0.05. At the meiosis stage, a total of 151 DEGs were identified, including 85 upregulated and 66 downregulated genes. By the young microspore stage, the number of DEGs increased to 194, comprising 134 upregulated and 60 downregulated transcripts. This expansion of the DEG population suggests that the absence of *RbohF*-mediated ROS signaling leads to a progressive divergence in gene expression patterns throughout microsporogenesis (**Fig. S7** and **Table S3-S5**).

### Validation of DEGs via RT-qPCR

To validate the reliability of the RNA-seq transcriptomic data, we performed RT-qPCR on several representative differentially expressed genes (DEGs). The expression patterns were examined at both the meiosis (Mei) and young microspore (YM) stages in the wild-type (T) and CRISPR *rbohf* (CF) lines. Consistent with the RNA-seq findings, the expression of *BURP15* (Os09g0480900) and the *C6 protein* gene (Os11g0582500) showed distinct stage-specific patterns, with *BURP15* being significantly downregulated in mutant lines during the Mei stage but showing robust expression at the YM stage (**Figure 10** **A, D; Figure S8 A, D**). In contrast, several defense- and drought-responsive genes were markedly upregulated in the CRISPR *rbohf* lines compared with the wild type. Specifically, two members of the GRAS transcription factor family, *OsGRAS23* (Os03g0629800 and Os01g0537250), which are known to mediate drought stress tolerance, exhibited a dramatic increase in expression in CF lines at both the Mei and YM stages (**Figure 10** **G, H, J, K; Figure S8 G, H, J, K**). Furthermore, the expression of *OsGLP1* (Os08g0460000), a germin-like protein involved in plant immunity and development, was also significantly higher in the mutant lines across both developmental stages (**Figure 10** **I, L; Figure S8 I, L**). Collectively, these RT-qPCR results highly correlate with the RNA-seq data, reinforcing the notion that the loss of *rbohF* function leads to the activation of specific stress and defense signaling pathways during pollen development.

**Fig. 10.**
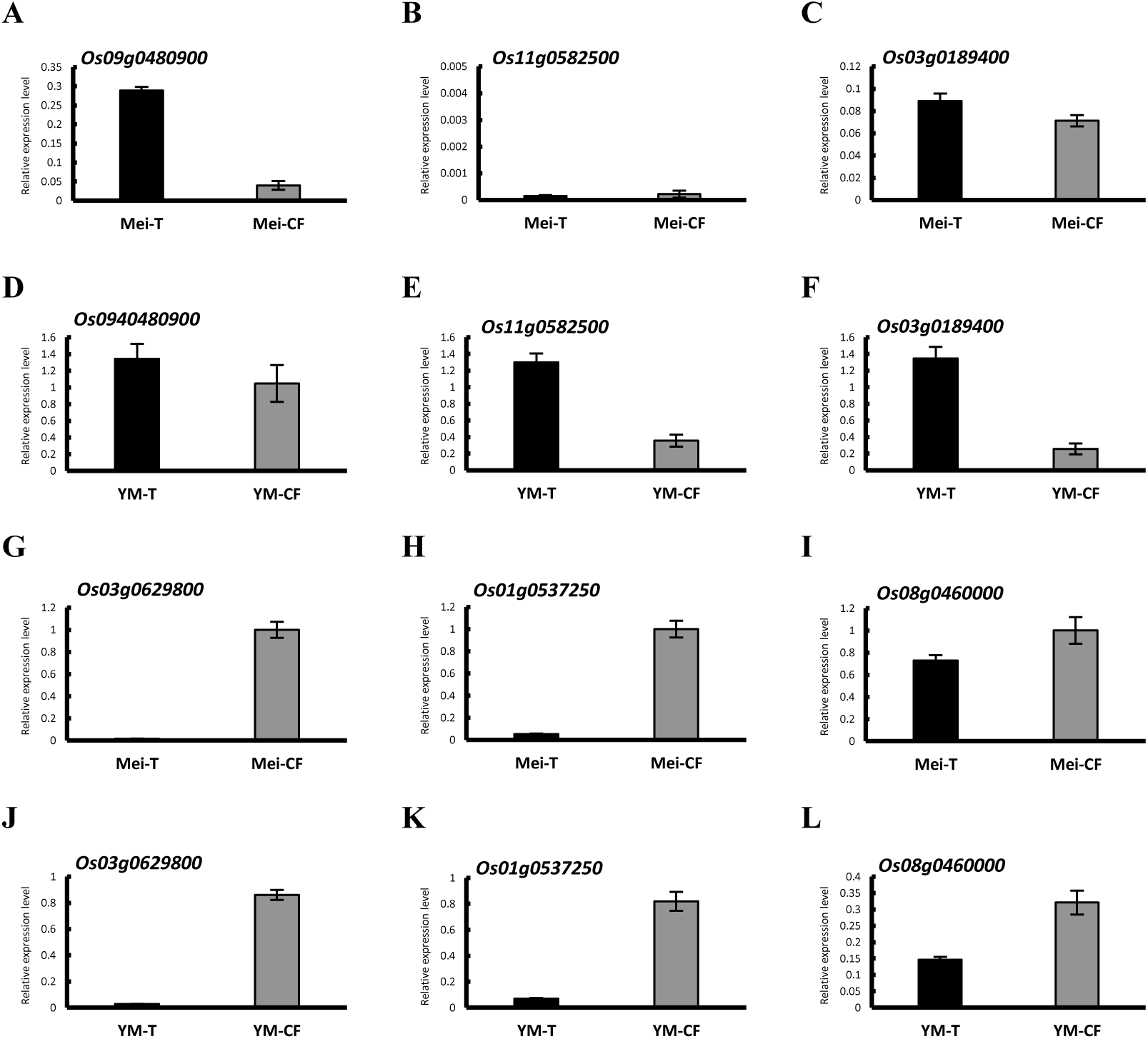
Expression patterns of significant DEGs at the meiosis (Mei) and young microspore (YM) stages in wild-type (T) and CRISPR *rbohf* (CF) lines. (A, B, C, G, H, I) Mei stage; (D, E, F, J, K, L) YM stage. Several stress- and defense-related genes were upregulated in the CRISPR *rbohf* lines. Os09g0480900 encodes BURP15. Os11g0582500 encodes the C6 protein. Os03g0629800 encodes OsGRAS23, a transcription factor that plays a key role in the response to drought stress. Os08g0460000 encodes OsGLP1, a germin-like protein involved in plant defense and developmental processes. Os01g0537250 also encodes OsGRAS23, which contributes to drought stress tolerance. Each bar represents the mean ± SEM of three biological replicates. P values were determined using an unpaired two-tailed Student’s t-test. Asterisks indicate statistically significant differences (*p* < 0.05) between wild-type and CRISPR *rbohf* lines.

## Discussion

The precise temporal coordination of tapetal PCD is essential for successful pollen development, with ROS acting as key signaling molecules in this process. Our results demonstrate that *RbohF* plays a central role in regulating ROS homeostasis during rice anther development. Loss of *RbohF* led to a marked reduction and delay in the accumulation of superoxide (OL•L) and hydrogen peroxide (H_2_O_2_) in the tapetum, particularly from the meiosis to young microspore (YM) stages (**Fig. 7; Supplementary Fig. S4**). These altered ROS dynamics were closely associated with delayed tapetal PCD, as evidenced by weakened and delayed TUNEL signals, ultimately resulting in defective pollen maturation and microspore collapse (**Fig. 5, 8**). These findings support the view that *RbohF*-mediated ROS production functions as a critical developmental signal required for timely tapetal degeneration.

The role of NADPH oxidase-derived ROS in anther development appears to be evolutionarily conserved. In *Arabidopsis*, AtRbohE has been shown to generate ROS required for tapetal PCD, and mutants exhibit pollen defects similar to those observed in *rbohf*. In their study, Xie and colleagues continually highlighted the importance of precisely regulated spatiotemporal ROS accumulation for the initiation of the PCD program (Xie *et al*., 2014). Our findings extend this framework to rice and identify *RbohF* as a major source of ROS in monocot anthers.

Notably, key transcriptional regulators of pollen development, including *UDT1*, *GAMYB*, *bHLH142/TIP2*, *TDR*, and *EAT1*, are essential for tapetal PCD, and loss of function of any of these factors results in defective PCD and aborted pollen development (Aya *et al*., 2009; Fu *et al*., 2014; Jung *et al*., 2005; Ko *et al*., 2014; Li *et al*., 2006; Niu *et al*., 2013). Our studies further showed that mutation of these transcription factors leads to significant downregulation of *RbohF* expression (**Fig. 9**). In contrast, the expression levels of these four transcription factors were largely unchanged in the *rbohf* mutant (**Supplementary Fig. S5**). Together, these results indicate that the *rbohf* mutant does not perturb the upstream transcriptional network; rather, *RbohF* functions downstream of, or in parallel with, these key regulators and primarily acts as an executor of ROS-dependent signaling during tapetal PCD.

Spatial and temporal control of ROS accumulation is critical for proper tapetal function. In wild-type anthers, ROS signals were predominantly localized to the tapetal layer and vascular bundle and peaked at the meiosis-to-YM stages. In contrast, *rbohf* mutants exhibited delayed and attenuated ROS accumulation, indicating that both the timing and intensity of ROS production are crucial. Insufficient ROS levels likely fail to activate downstream proteases and nucleases required for PCD execution, resulting in prolonged tapetal persistence and impaired nutrient supply to developing microspores.

In addition to its developmental role, *RbohF*-mediated ROS signaling contributes to environmental adaptability during the reproductive stages. The two cropping seasons analyzed differed in temperature profiles, with the 2024-I season experiencing cooler conditions (**Fig. 3,4; Supplementary Fig. S3**). Rice is particularly sensitive to low temperatures during the YM stage, where temperatures below 17°C disrupt tapetal PCD and induce pollen sterility. Previous studies have shown that cool temperatures delay tapetal degradation and impair carbohydrate metabolism, partly through repression of *OsINV4* (Gothandam *et al*., 2007; Oda *et al*., 2010; Oliver *et al*., 2005). Consistent with these findings, *rbohf* mutants exhibited more severe sterility during the cooler 2024-I cropping season, when the minimum temperature reached 15.7°C during the critical pollen developmental stage. This increased sterility was accompanied by a pronounced reduction in reactive oxygen species (ROS) accumulation, as well as a significant decline in pollen viability compared with the wild-type plants. These observations indicate that RbohF-derived ROS likely play a protective and regulatory role under suboptimal temperature conditions. Specifically, ROS generated by RbohF may function as a buffering or stabilizing signal that helps sustain normal pollen development, possibly by modulating redox homeostasis, maintaining cellular integrity, or supporting temperature-sensitive metabolic and signaling pathways. In the absence of adequate RbohF activity, this buffering capacity appears compromised, leading to disrupted developmental progression and increased susceptibility to cold-induced reproductive failure.

Although *RbohF* appears to play a major role in pollen development, the knockout mutant exhibited only partial sterility (**Fig. 3,4**). This incomplete phenotype suggests potential functional redundancy within the rice *Rboh* gene family. Presumably, other *Rboh* members may partially compensate by contributing to ROS production during anther development, thereby mitigating the impact of *RbohF* loss. The rice genome encodes multiple Rboh isoforms with potentially distinct expression patterns and functions. Evidence from *Arabidopsis* indicates that different Rboh proteins, such as AtRbohD and AtRbohE, can act redundantly or in a tissue-specific manner (Kwak *et al*., 2003). Therefore, it is plausible that other rice Rboh members function in parallel with *RbohF* or partially compensate for its loss. Further investigation using higher-order mutants and spatial expression analyses will be necessary to clarify the extent of functional redundancy and specialization within this gene family.

In conclusion, *RbohF* is essential for maintaining ROS homeostasis during rice anther development. Its loss disrupts the timely accumulation of ROS, leading to delayed tapetal PCD and defective pollen maturation. The increased sensitivity of *rbohf* mutants to cool temperatures further highlights the role of *RbohF* in integrating developmental and environmental signals. Together, these findings establish *RbohF* as a key regulator of reproductive development and stress resilience in rice.

## Acknowledgements

We thank the Plant Technology Core Facility at Academia Sinica for generating the CRISPR/Cas9-edited lines. We are grateful to the Transgenic Plant Laboratory at Academia Sinica for rice transformation. We also thank the GMO greenhouse at the Biotechnology Center in Southern Taiwan (AS-BCST) and RNA-ISH Core for providing growth space and technical support. Confocal microscopy was performed at the Confocal Microscope Core Facility of AS-BCST, and DNA sequencing services were provided by the DNA Sequencing Core Facility at the Institute of Biomedical Sciences, Academia Sinica. We thank Miranda Loney for English editing. This work was supported by the Ministry of Science and Technology (Taiwan) to S.-S. Ko (MOST 111-2313-B-001-009-MY3). We gratefully acknowledge MOST for supporting the postdoctoral grant awarded to H.-C. Lu (MOST- 114-2811-B-001-037).

## Author contributions

S.-S.K. conceived and designed the study. H.-C.L. and S.-S.K. wrote the manuscript. M.-J.L. verified the CRISPR-induced mutations and performed RT–qPCR analyses. H.-C.L. conducted phylogenetic and RNA-seq analyses. J.-C.H. and H.-C.H. carried out phenotypic analyses and ROS staining. T.-T.Y. performed *in situ* hybridization and TUNEL assays. Y.-P.H. conducted tissue sectioning.

## Data Availability

The RNA-seq datasets generated and analyzed in the current study are available in the NCBI BioProject repository, https://www.ncbi.nlm.nih.gov/bioproject/PRJNA1445505.

---

**Supplementary Table S1.**
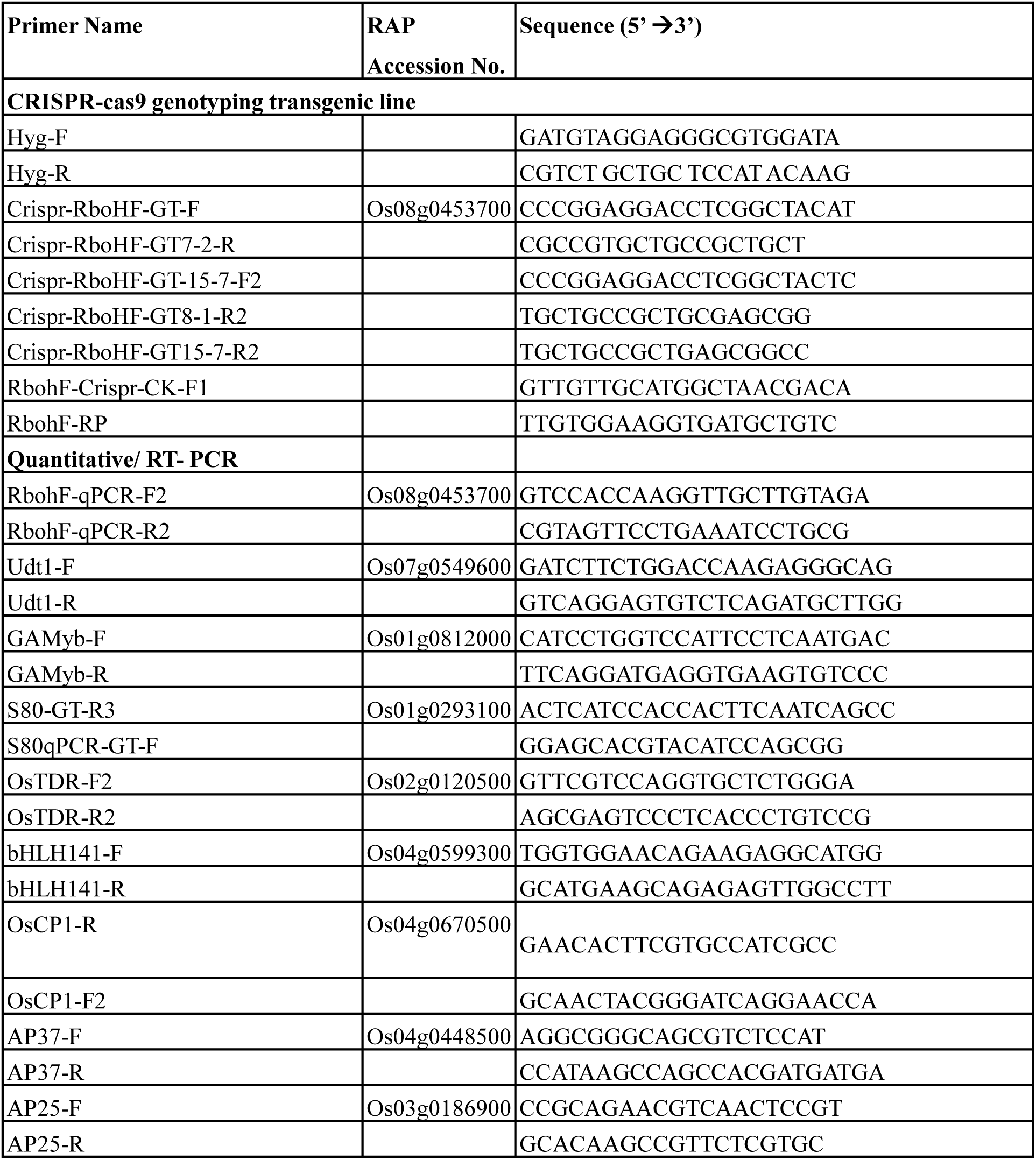
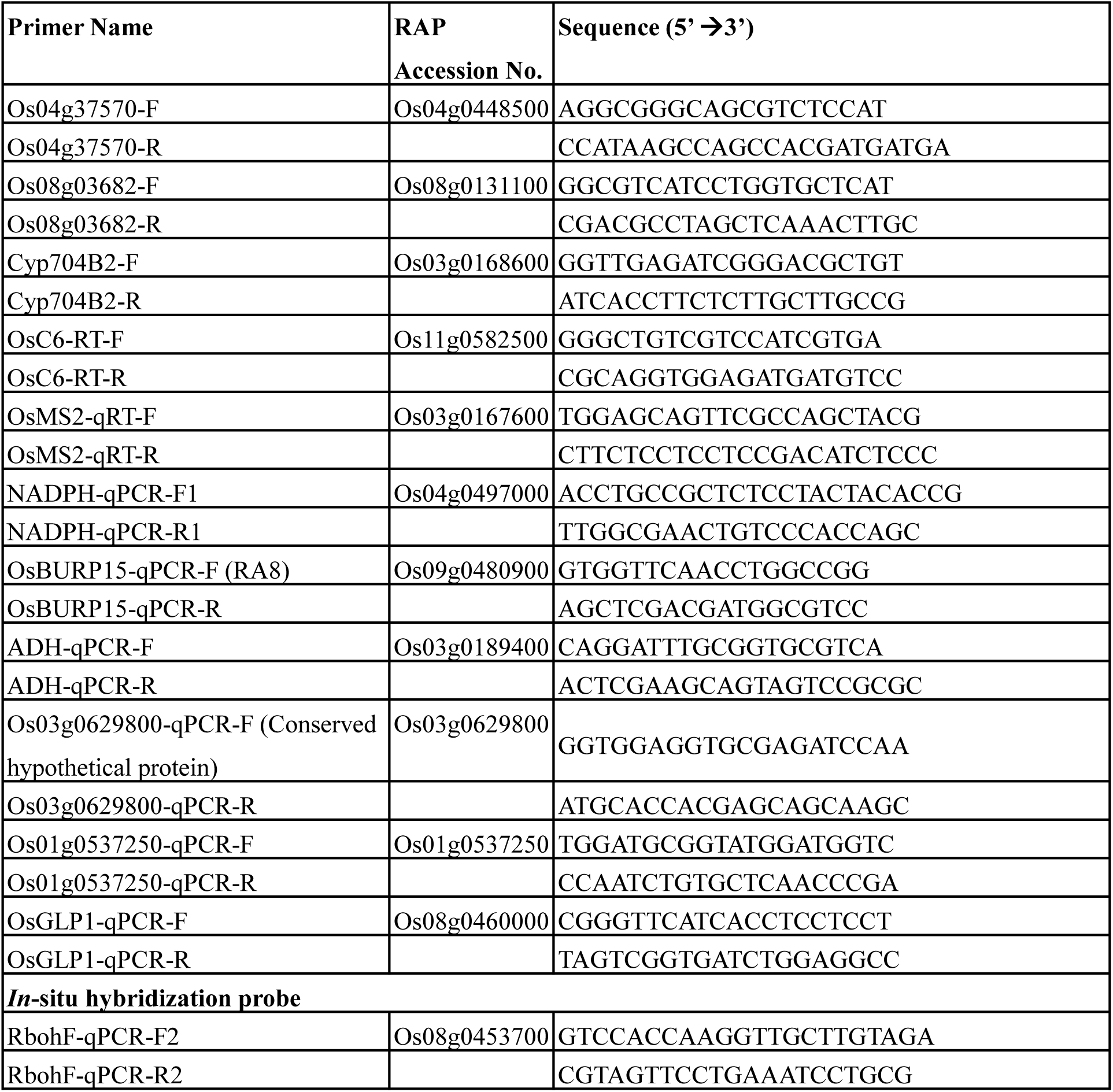
List of primers used in this study.

**Supplementary Table S2.** Top 20 DEGs significantly upregulated in *rbohf* mutant compared to TNG67 during meiotic stages.

| geneID | log2FoldChange | pvalue | Description |
| --- | --- | --- | --- |
| Os01g0537250 | 12.2332721 | 3.69962E-32 | Protein of unknown function DUF3778 domain containing protein. (Os01t0537250-01) |
| Os07g0442800 | 10.20358625 | 2.31122E-22 | Conserved hypothetical protein. (Os07t0442800-01) |
| Os03g0115800 | 10.02775399 | 5.3405E-175 | Conserved hypothetical protein. (Os03t0115800-01)%3BConserved hypothetical protein. (Os03t0115800-02) |
| Os09g0463800 | 9.868961533 | 5.41006E-16 | Conserved hypothetical protein. (Os09t0463800-01) |
| Os10g0474600 | 8.498571862 | 1.05106E-05 | Conserved hypothetical protein. (Os10t0474600-00) |
| Os06g0685400 | 7.939769402 | 1.53271E-10 | Conserved hypothetical protein. (Os06t0685400-01) |
| Os03g0105500 | 7.676217454 | 3.07922E-13 | Conserved hypothetical protein. (Os03t0105500-01) |
| Os02g0157000 | 7.112671165 | 2.34355E-08 | Leucine-rich repeat domain containing protein. (Os02t0157000-00) |
| Os01g0262401 | 6.955042217 | 1.87786E-10 | Hypothetical genes. (Os01t0262401-00) |
| Os06g0295000 | 6.603134307 | 7.85175E-07 | Hypothetical conserved gene. (Os06t0295000-01) |
| Os03g0629800 | 6.484626297 | 8.70405E-09 | Conserved hypothetical protein. (Os03t0629800-01) |
| Os05g0580200 | 6.483338281 | 6.51572E-07 | Conserved hypothetical protein. (Os05t0580200-01) |
| Os12g0555000 | 6.359021741 | 3.61739E-06 | Similar to Probenazole-inducible protein PBZ1. (Os12t0555000-01) |
| Os07g0538000 | 6.143345961 | 1.25974E-05 | Similar to hydrolase%2C hydrolyzing O-glycosyl compounds. (Os07t0538000-01) |
| Os05g0524500 | 5.93272266 | 8.68065E-05 | Protein kinase%2C core domain containing protein. (Os05t0524500-01) |
| Os07g0122100 | 5.897416559 | 0.000223514 | Protein of unknown function DUF1719%2C Oryza sativa family protein. (Os07t0122100-01) |
| Os10g0432200 | 5.814278009 | 3.49484E-34 | Conserved hypothetical protein. (Os10t0432200-01) |
| Os06g0559600 | 5.628388231 | 5.06578E-05 | Hypothetical conserved gene. (Os06t0559600-01) |
| Os11g0278900 | 5.524196303 | 8.43917E-05 | Conserved hypothetical protein. (Os11t0278900-01) |
| Os04g0339800 | 5.248091545 | 5.18092E-06 | Similar to predicted protein. (Os04t0339800-00) |

**Supplementary Table S3.**
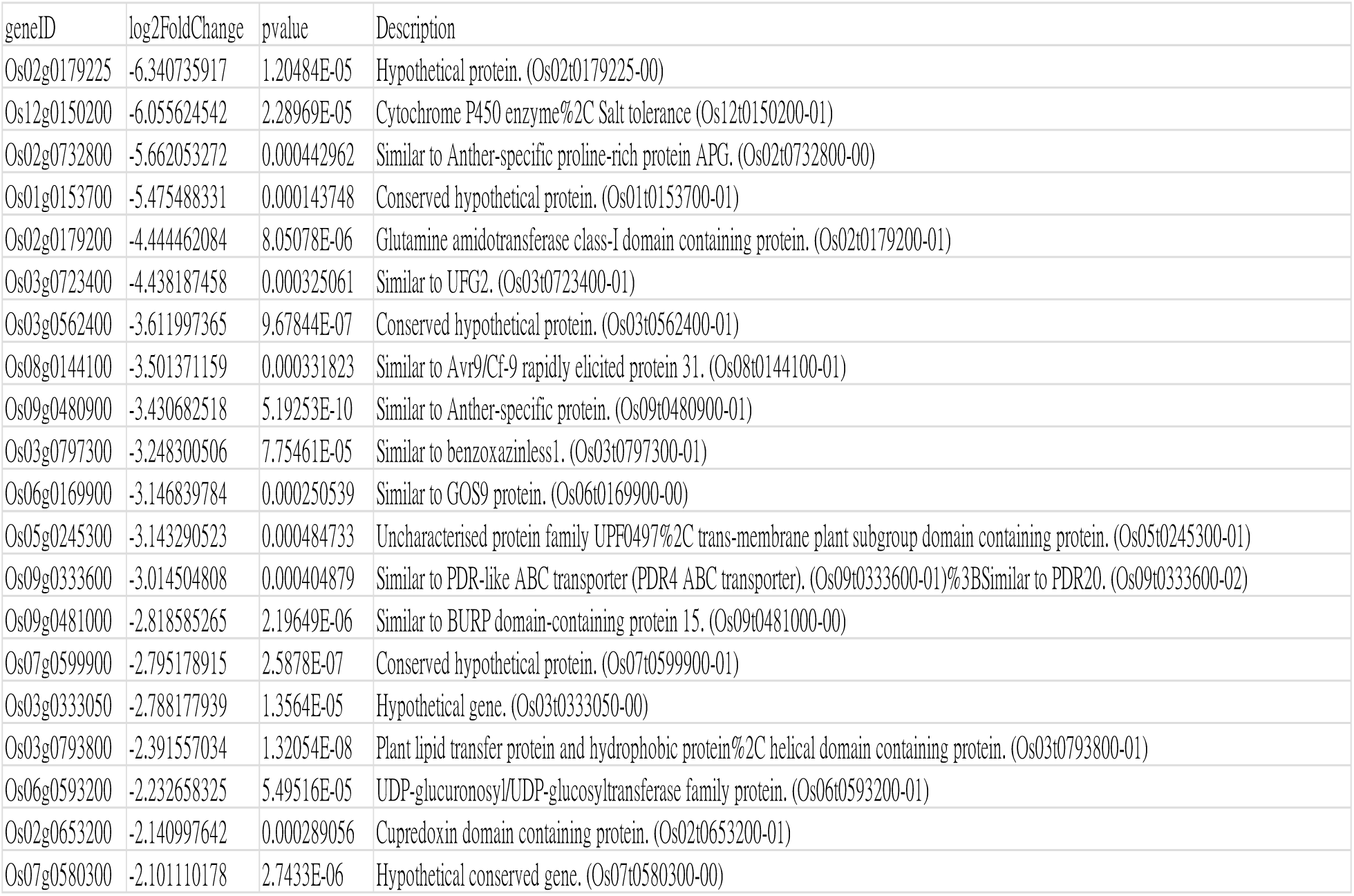
Top 20 DEGs significantly downregulated in *rbohf* compared to TNG67 during meiotic stages.

**Supplementary Table S4.** Top 20 DEGs significantly upregulated in *rbohf* compared to TNG67 during YM stages.

| geneID | log2FoldChange | pvalue | Description |
| --- | --- | --- | --- |
| Os07g0442800 | 11.95968619 | 7.76385E-24 | Conserved hypothetical protein. (Os07t0442800-01) |
| Os03g0115800 | 10.40913574 | 1.0724E-149 | Conserved hypothetical protein. (Os03t0115800-01)%3BConserved hypothetical protein. (Os03t0115800-02) |
| Os09g0463800 | 10.05961476 | 1.042E-16 | Conserved hypothetical protein. (Os09t0463800-01) |
| Os01g0537250 | 10.03591453 | 5.60794E-72 | Protein of unknown function DUF3778 domain containing protein. (Os01t0537250-01) |
| Os12g0555000 | 8.786752182 | 4.18962E-13 | Similar to Probenazole-inducible protein PBZ1. (Os12t0555000-01) |
| Os01g0262401 | 7.713562506 | 1.99008E-13 | Hypothetical genes. (Os01t0262401-00) |
| Os06g0685400 | 7.70238282 | 5.67026E-10 | Conserved hypothetical protein. (Os06t0685400-01) |
| Os06g0295000 | 7.14951272 | 1.53179E-08 | Hypothetical conserved gene. (Os06t0295000-01) |
| Os07g0511301 | 6.75079624 | 5.36152E-10 | Hypothetical protein. (Os07t0511301-00) |
| Os09g0344500 | 6.565636807 | 9.08758E-07 | Similar to O-methyltransferase ZRP4 (EC 2.1.1.-) (OMT). (Os09t0344500-01) |
| Os05g0580200 | 6.447489257 | 1.66475E-06 | Conserved hypothetical protein. (Os05t0580200-01) |
| Os07g0511400 | 6.417976954 | 5.4627E-127 | Hypothetical protein. (Os07t0511400-01) |
| Os11g0236932 | 6.320699846 | 3.16701E-06 | Hypothetical conserved gene. (Os11t0236932-01) |
| Os06g0662850 | 6.097933689 | 1.00786E-05 | Similar to OSIGBa0113K06.3 protein. (Os06t0662850-00) |
| Os01g0387566 | 6.067743057 | 7.44918E-06 | Conserved hypothetical protein. (Os01t0387566-00) |
| Os10g0432200 | 5.788744016 | 1.6967E-165 | Conserved hypothetical protein. (Os10t0432200-01) |
| Os11g0278900 | 5.705101019 | 3.54497E-05 | Conserved hypothetical protein. (Os11t0278900-01) |
| Os03g0619850 | 5.623770721 | 0.000121705 | Putative B3 domain-containing protein Os03g0619850. (Os03t0619850-00) |
| Os07g0538000 | 5.601359635 | 8.11337E-05 | Similar to hydrolase%2C hydrolyzing O-glycosyl compounds. (Os07t0538000-01) |
| Os08g0203350 | 5.54452394 | 8.23554E-10 | Conserved hypothetical protein. (Os08t0203350-01) |

**Supplementary Table S5.** Top 20 DEGs significantly downregulated in *rbohf* compared to TNG67 during YM stages.

| geneID | log2FoldChange | pvalue | Description |
| --- | --- | --- | --- |
| Os02g0179225 | -6.340735917 | 1.20484E-05 | Hypothetical protein. (Os02t0179225-00) |
| Os12g0150200 | -6.055624542 | 2.28969E-05 | Cytochrome P450 enzyme%2C Salt tolerance (Os12t0150200-01) |
| Os02g0732800 | -5.662053272 | 0.000442962 | Similar to Anther-specific proline-rich protein APG. (Os02t0732800-00) |
| Os01g0153700 | -5.475488331 | 0.000143748 | Conserved hypothetical protein. (Os01t0153700-01) |
| Os02g0179200 | -4.444462084 | 8.05078E-06 | Glutamine amidotransferase class-I domain containing protein. (Os02t0179200-01) |
| Os03g0723400 | -4.438187458 | 0.000325061 | Similar to UFG2. (Os03t0723400-01) |
| Os03g0562400 | -3.611997365 | 9.67844E-07 | Conserved hypothetical protein. (Os03t0562400-01) |
| Os08g0144100 | -3.501371159 | 0.000331823 | Similar to Avr9/Cf-9 rapidly elicited protein 31. (Os08t0144100-01) |
| Os09g0480900 | -3.430682518 | 5.19253E-10 | Similar to Anther-specific protein. (Os09t0480900-01) |
| Os03g0797300 | -3.248300506 | 7.75461E-05 | Similar to benzoxazinless1. (Os03t0797300-01) |
| Os06g0169900 | -3.146839784 | 0.000250539 | Similar to GOS9 protein. (Os06t0169900-00) |
| Os05g0245300 | -3.143290523 | 0.000484733 | Uncharacterised protein family UPF0497%2C trans-membrane plant subgroup domain containing protein. (Os05t0245300-01) |
| Os09g0333600 | -3.014504808 | 0.000404879 | Similar to PDR-like ABC transporter (PDR4 ABC transporter). (Os09t0333600-01)%3BSimilar to PDR20. (Os09t0333600-02) |
| Os09g0481000 | -2.818585265 | 2.19649E-06 | Similar to BURP domain-containing protein 15. (Os09t0481000-00) |
| Os07g0599900 | -2.795178915 | 2.5878E-07 | Conserved hypothetical protein. (Os07t0599900-01) |
| Os03g0333050 | -2.788177939 | 1.3564E-05 | Hypothetical gene. (Os03t0333050-00) |
| Os03g0793800 | -2.391557034 | 1.32054E-08 | Plant lipid transfer protein and hydrophobic protein%2C helical domain containing protein. (Os03t0793800-01) |
| Os06g0593200 | -2.232658325 | 5.49516E-05 | UDP-glucuronosyl/UDP-glucosyltransferase family protein. (Os06t0593200-01) |
| Os02g0653200 | -2.140997642 | 0.000289056 | Cupredoxin domain containing protein. (Os02t0653200-01) |
| Os07g0580300 | -2.101110178 | 2.7433E-06 | Hypothetical conserved gene. (Os07t0580300-00) |

**Supplementary Fig. S1.**
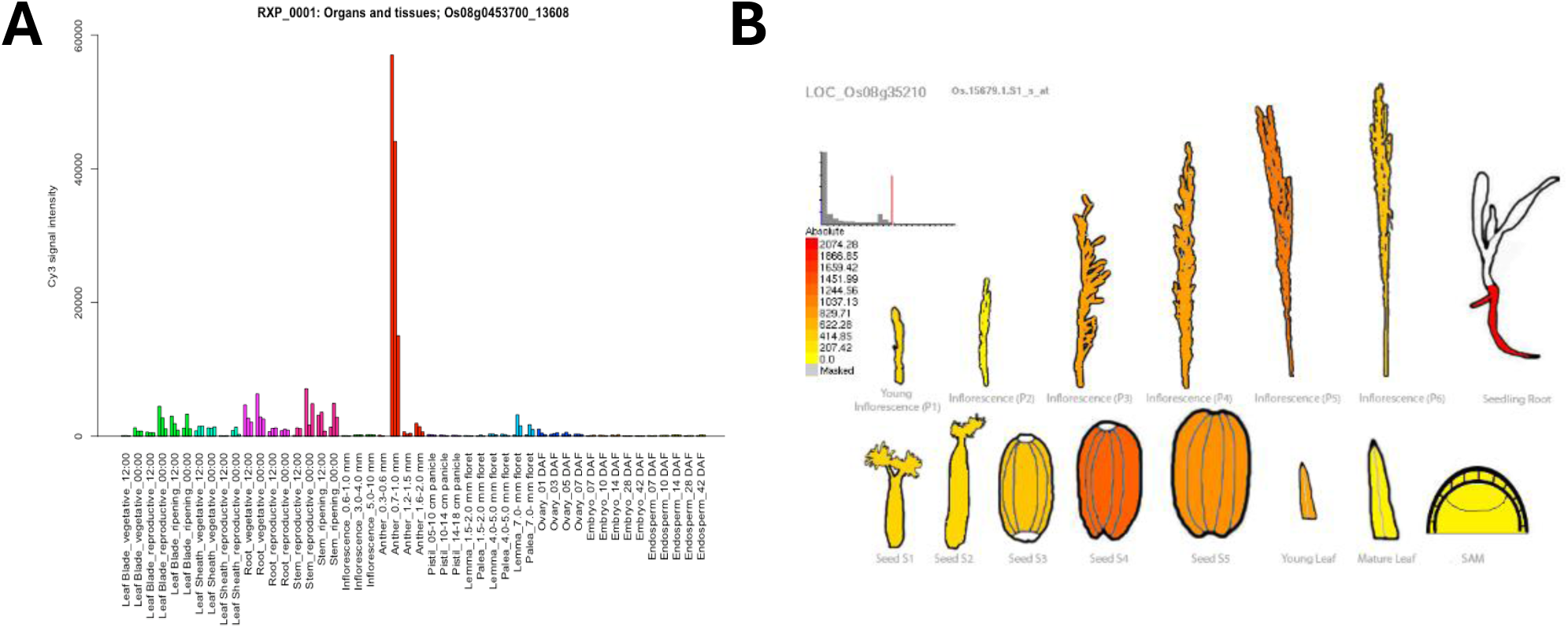
Gene expression patterns of *RbohF* in rice. (A) gene locus Os08g0453700 (LOC_Os08g35210) in the RiceXPro database and (B) Rice eFP browser.

**Supplementary Fig. S2.**
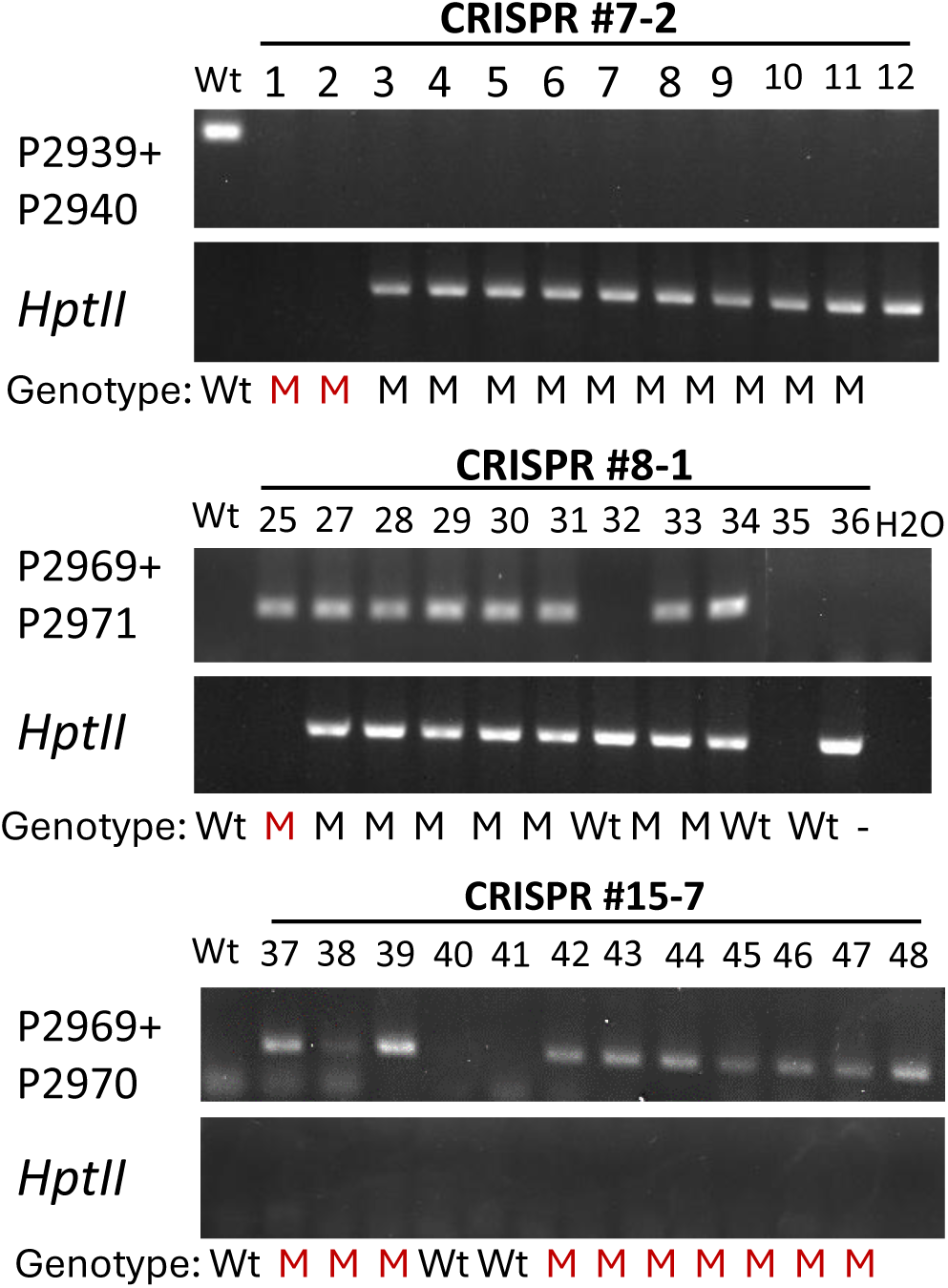
Genomic PCR validation of CRISPR-mediated *rbohf* mutants. Representative gel images show the amplification of target gene regions (upper panels) and the HptII selection marker (lower panels). Specific primer were employed to detect sequence variations at the target loci. Wt represents the wild-type allele, while M indicates the presence of a mutant allele.

**Supplementary Fig. S3.**
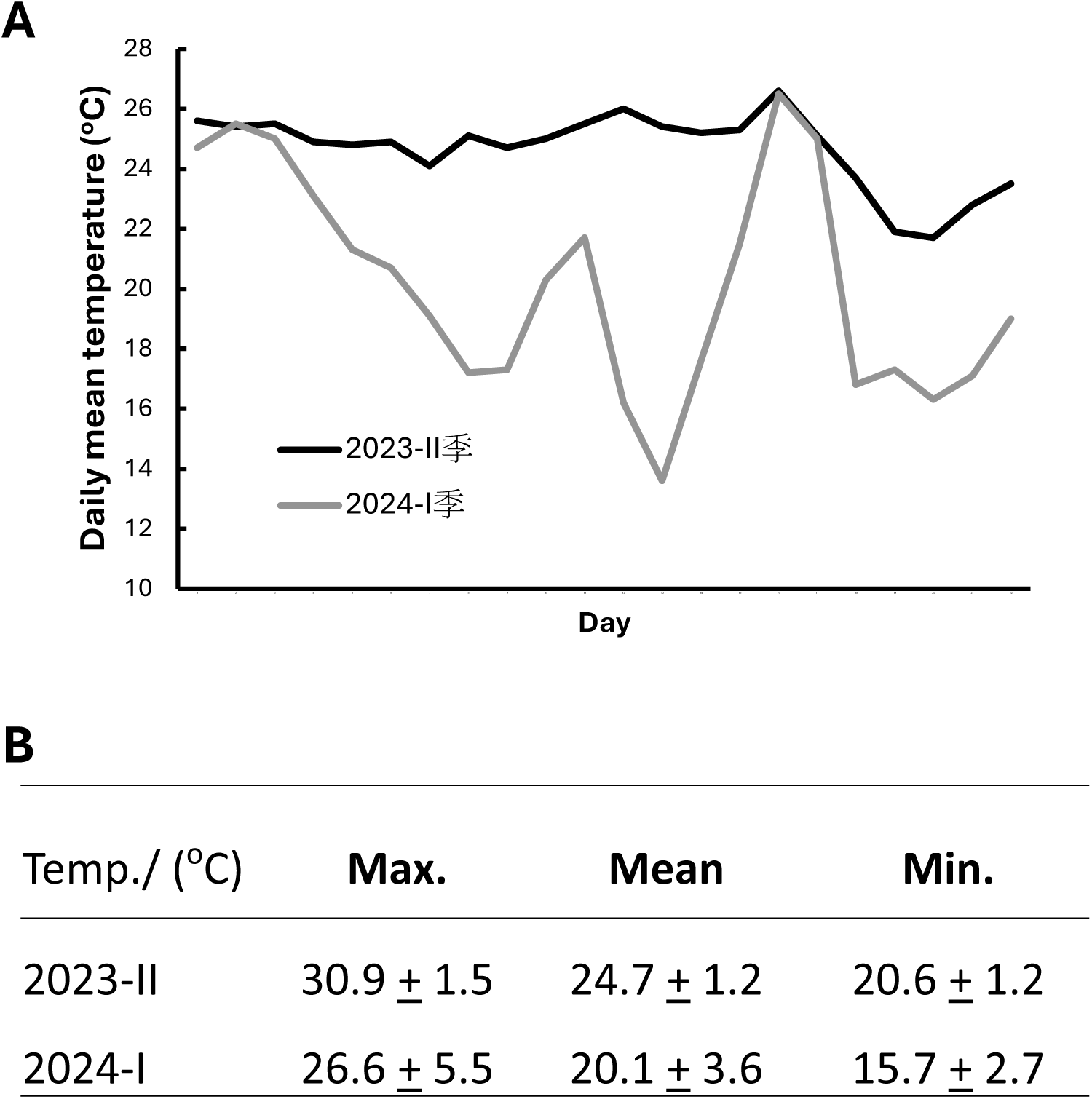
Temperature records during rice pollen developmental stages in the second cropping season of 2023 and the first cropping season of 2024.

**Supplementary Fig. S4.**
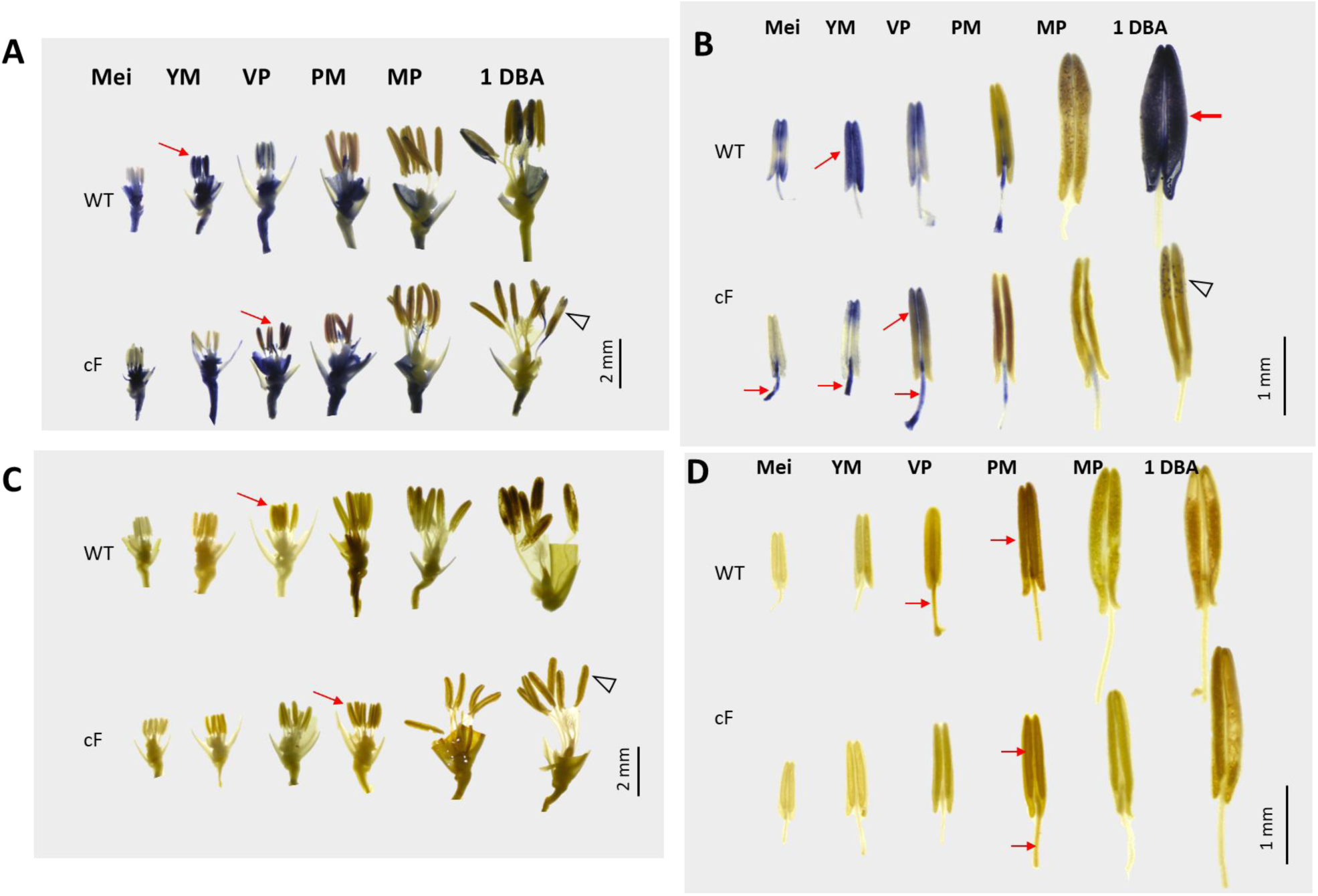
ROS distribution in the developing anthers of the WT (TNG67) and CRISPR *rbohf* (cF) transgenic line #7-2. (A,B) Developing spikelets and anthers stained with superoxide by using NBT. (C, D) stained H_2_O_2_ by DAB. Arrows showed ROS signal. Arrow heads showed weak ROS. open triangles mark regions with delayed ROS production Bar, 2 mm (A, C), 1mm (B, D).

**Supplementary Fig. S5.**
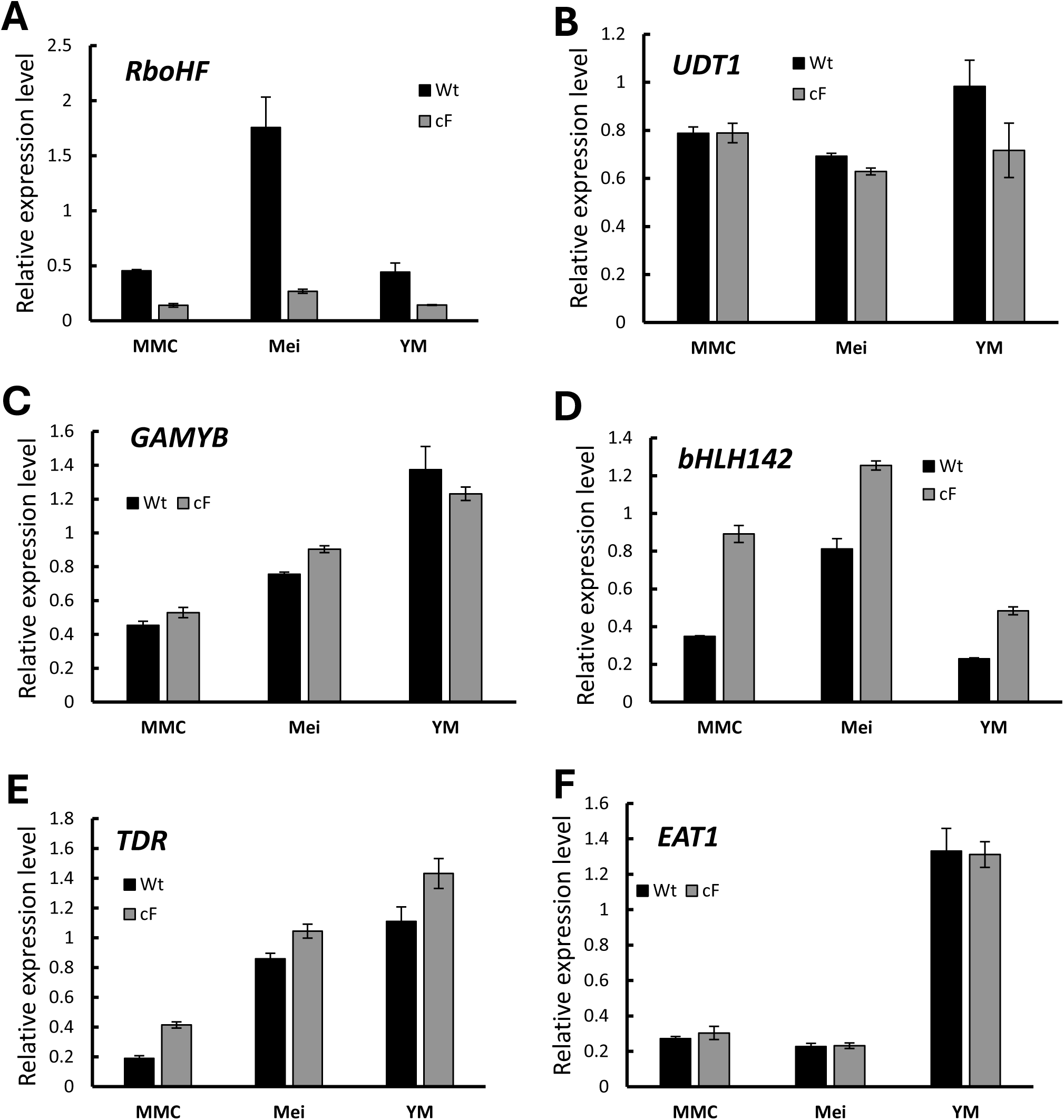
CRISPR-Cas9-mediated knockout of *RbohF* modifies the expression of key transcription factors involved in pollen development.

**Supplementary Fig. S6.**
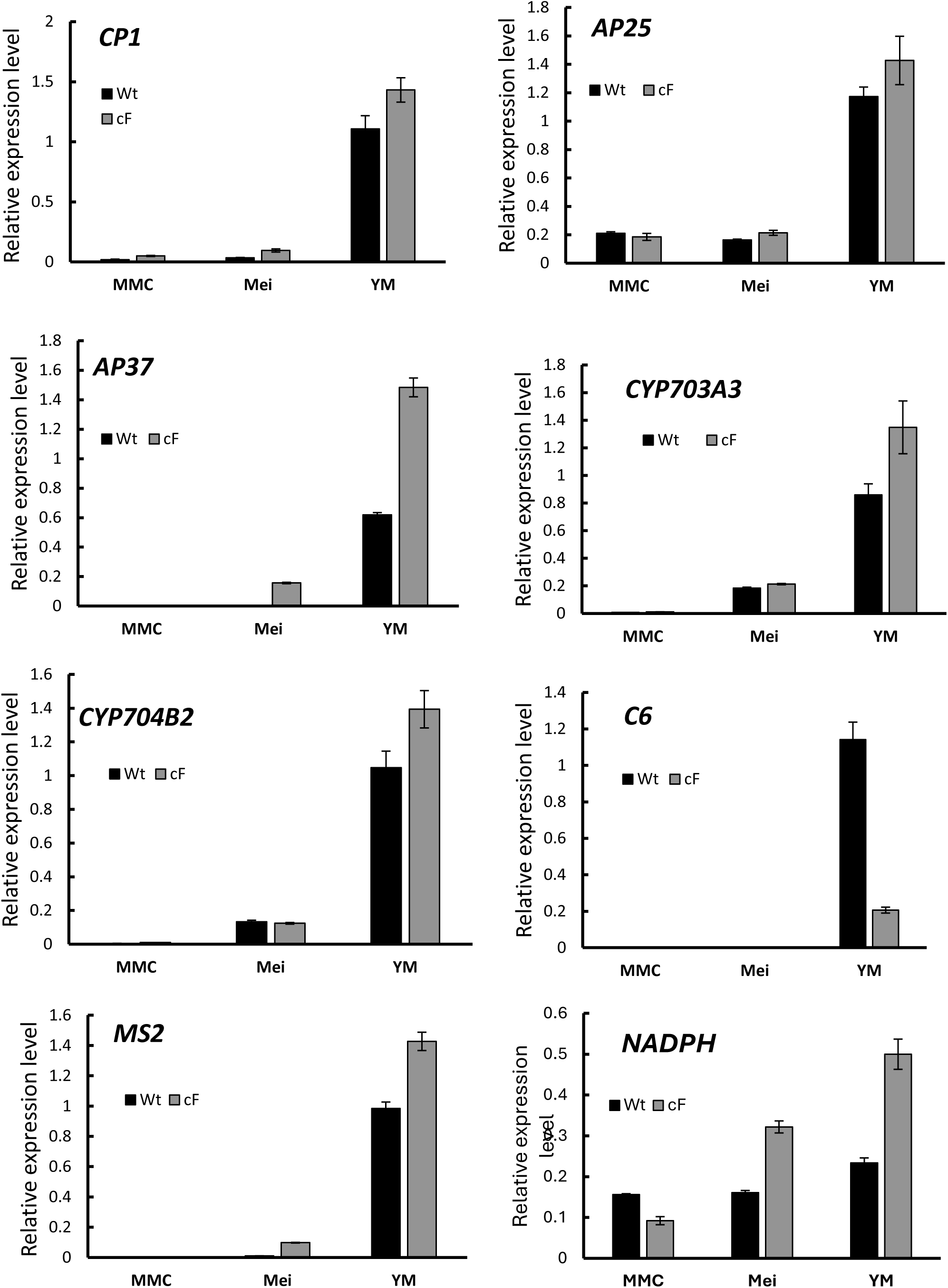
RT-qPCR analysis showing that CRISPR *rbohF* mutation alters the expression patterns of pollen marker genes. Anthers at the microspore mother cell (MMC), meiosis (Mei), and young microspore (YM) stages from wild-type (WT) plants and the CRISPR *rbohF* mutant were collected for RNA isolation and RT-qPCR analysis. Data represent tbree replicates.

**Supplementary Fig. S7.**
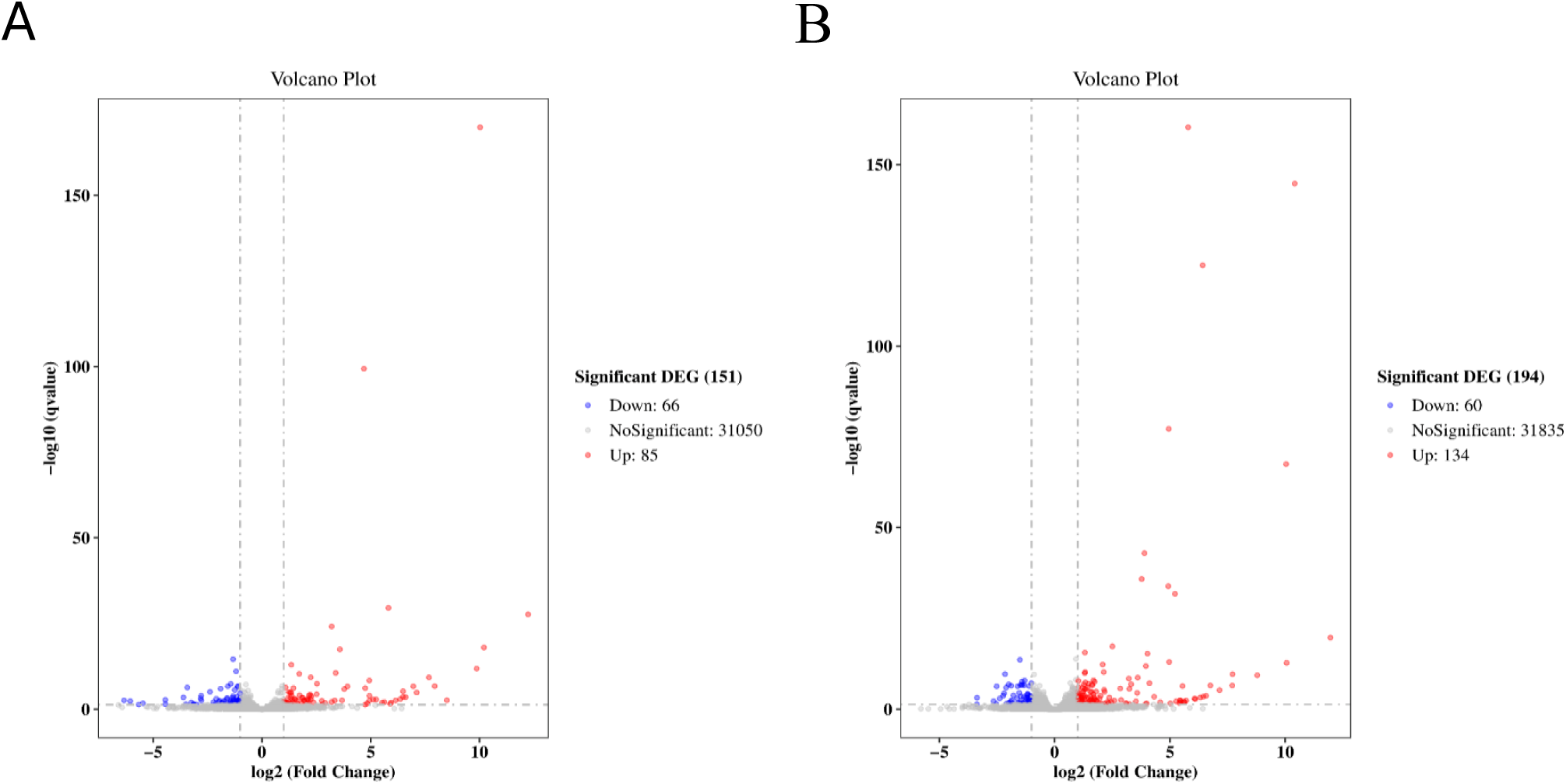
A volcano plot of DEGs (FC > 2 and adjusted *p* < 0.05) at the Mei (A) and YM stage (B). The horizontal axis represents the fold change, and the vertical axis represents the adjusted p-value. The red and blue circles indicate up- and down-regulated genes, respectively.

**Supplementary Fig. S8.**
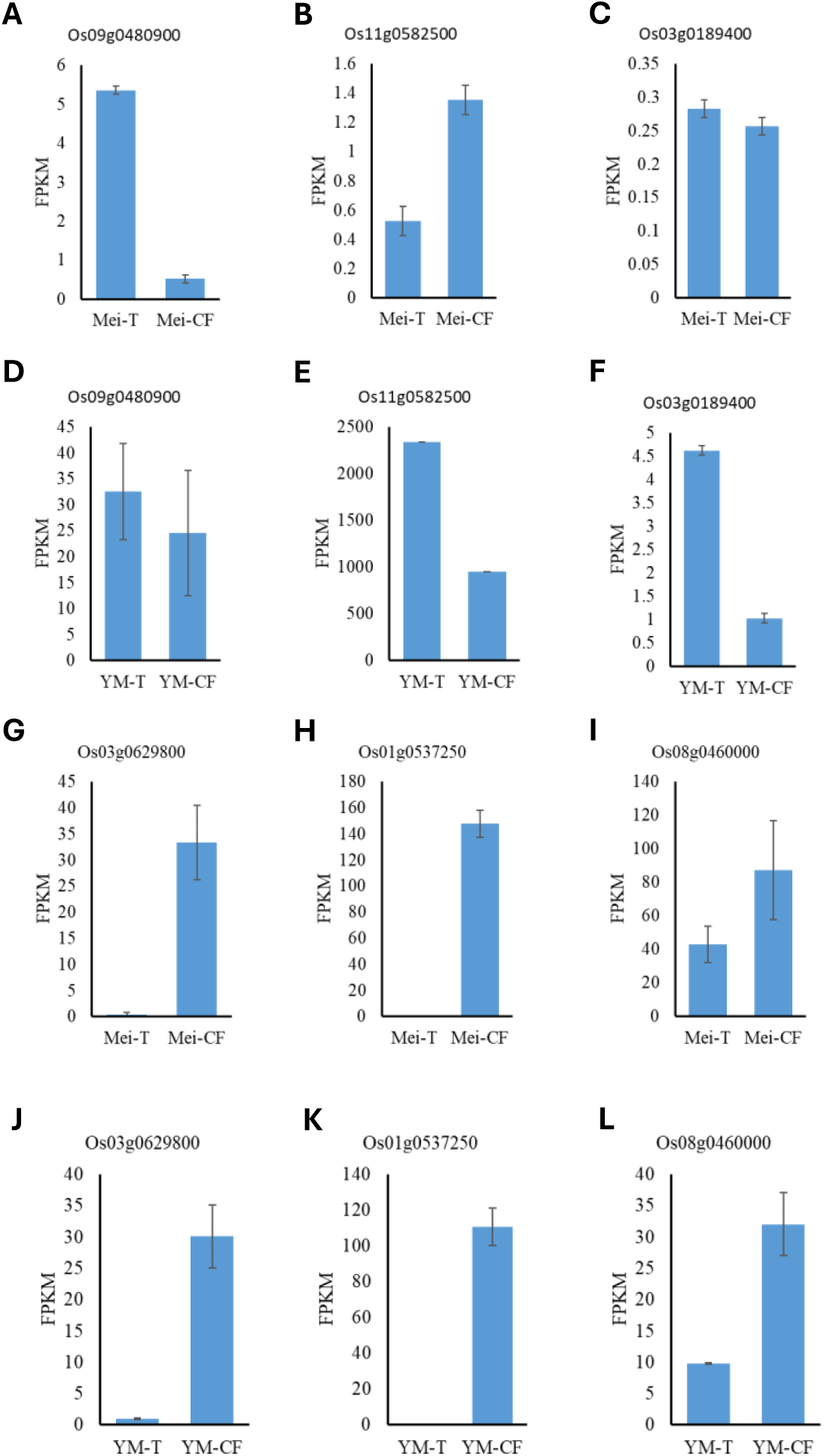
Expression patterns of significantly DEGs at the meiosis (Mei) and young microspore (YM) stages in wild-type (T) and CRISPR *rbohF* (CF) lines. (A, B, C, G, H, I) Mei stage; (D, E, F, J, K, L) YM stage. Several stress- and defense-related genes were notably upregulated in the CRISPR *rbohF* lines. Os09g0480900 encodes BURP15. Os11g0582500 encode C6 protein. *Os03g0629800* encodes OsGRAS23, a transcription factor that plays a key role in the response to drought stress. *Os08g0460000* encodes OsGLP1, a germin-like protein involved in plant defense and developmental processes. *Os01g0537250* also encodes OsGRAS23, which contributes to drought stress tolerance. Each bar represents the mean ± SEM of three biological replicates.

